# Nuclear eNOS interacts with and *S*-nitrosates ADAR1 to modulate type I interferon signaling and endothelial function

**DOI:** 10.1101/2024.05.24.595750

**Authors:** Xiaozhu Zhou, Carsten Kuenne, Stefan Günther, Ilka Wittig, Beyza Güven, Doha Boutguetait, Fredy Delgado Lagos, Nadja Sachs, Lars Maegdefessel, Christian Münch, Stefan Offermanns, Ingrid Fleming, Mauro Siragusa

**Affiliations:** Goethe University, Institute for Vascular Signalling, Centre for Molecular Medicine, Frankfurt am Main, Germany; Max Planck Institute for Heart and Lung Research, Bioinformatics and Deep Sequencing Platform, Bad Nauheim, Germany; Goethe University, Functional Proteomics, Institute of Cardiovascular Physiology, Frankfurt am Main, Germany; Goethe University, Institute of Molecular Systems Medicine, Faculty of Medicine, Frankfurt am Main, Germany; Technical University Munich, Department for Vascular and Endovascular Surgery, Klinikum rechts der Isar, Germany; Technical University Munich, Institute of Molecular Vascular Medicine, Germany; Karolinska Institutet and University Hospital, Department of Medicine, Stockholm, Sweden; Max Planck Institute for Heart and Lung Research, Department of Pharmacology, Bad Nauheim, Germany; Goethe University, Centre for Molecular Medicine, Frankfurt am Main, Germany; German Center for Cardiovascular Research (DZHK), Partner site Munich Heart Alliance; German Center for Cardiovascular Research (DZHK), Partner site RheinMain, Frankfurt am Main and Bad Nauheim, Germany; CardioPulmonary Institute, Frankfurt am Main and Bad Nauheim, Germany

**Keywords:** eNOS, *S*-nitrosation, ADAR1, dsRNA, type IFN signaling

## Abstract

**Background:** Nitric oxide (NO) generated by the endothelial NO synthase (eNOS) regulates vascular tone and endothelial homeostasis to counteract vascular inflammation. Most eNOS is localized at the cell membrane or in the Golgi apparatus, but the enzyme is also present in the endothelial cell nucleus. Here we assessed the relevance of nuclear eNOS and NO signaling for endothelial cell function.

**Methods:** eNOS gain- and loss-of-function approaches were combined with confocal microscopy, biochemical, histological and multi-omics analyses. The pathophysiological relevance of the findings was assessed in murine models of atherogenesis (ApoE^-/-^ and partial carotid ligation) as well as in samples from patients with atherosclerosis.

**Results:** eNOS was present in the nucleus of unstimulated human and murine endothelial cells (*in vitro* and *ex vivo*) and stimulation with vascular endothelial growth factor (VEGF) enhanced its nuclear localization. Co-immunoprecipitation studies coupled with proteomics revealed the association of nuclear eNOS with 81 proteins involved in RNA binding and processing. Among the latter was double-stranded RNA-specific adenosine deaminase (ADAR1), an enzyme involved in editing double-stranded RNA (dsRNA) via the deamination of adenosine (A) to inosine (I). ADAR1 was *S*-nitrosated in human endothelial cells, and the knockdown of eNOS resulted in altered ADAR1-mediated A-to-I editing and an increase in dsRNA. Mitochondrial antiviral signaling protein (MAVS) was the transcript most affected by eNOS depletion, and endothelial cells lacking eNOS accumulated dsRNA as well as MAVS protein aggregates. These changes resulted in activation of the type I interferon (IFN) signaling pathway and a marked downregulation of cell cycle-related genes. ADAR1 depletion elicited similar effects on the activation of type I IFN signaling. As a result, growth factor-stimulated cell proliferation was abrogated and basal as well as stimulated cell death were increased. Endothelial dysfunction in mice as well as in subjects with atherosclerosis was accompanied by the accumulation of dsRNA and the activation of the type I IFN signaling. Preserving NO bioavailability *in vivo* in an eNOS gain-of-function mouse model (Tyr657Phe eNOS) prevented these effects.

**Conclusions:** Our findings have uncovered a novel mechanism linking nuclear eNOS-derived NO with the activity of ADAR1 to maintain vascular homeostasis. Reduced NO bioavailability results in the previously unrecognized activation of a type I IFN response in the endothelium, which contributes to atherogenesis.

## Introduction

Nitric oxide (NO) is a gasotransmitter with well-documented effects on vascular tone and endothelial homeostasis. In the endothelium, NO is generated by the endothelial NO synthase (eNOS), and its dysregulation has been linked with the development of cardiovascular disease.^1^ The effects of NO are attributed to its interaction with heme-containing proteins, such as soluble guanylyl cyclase and to the post-translational modification i.e. *S*-nitrosation, of susceptible cysteine residues in target proteins. Early work on the transcription factor nuclear factor κB demonstrated the impact of NO on gene expression.^2^ Since then a wealth of evidence has accumulated that links endogenously generated as well as exogenously applied NO with the post-translational modification of histones as well as DNA methylation and changes in microRNAs.^3^ Indeed, in tumor cells, the differential expression of thousands of transcripts in response to NO was associated with the post-translational modification of histones.^3^

The subcellular compartmentalization of eNOS is crucial for spatial NO production and signal transduction. In endothelial cells, the majority of eNOS is found in the perinuclear Golgi apparatus^4^ and in caveolae.^5^ However, the enzyme is also present in the endothelial cell nucleus and several stimuli e.g. vascular endothelial growth factor (VEGF),^6^ estrogen,^7^ and lysophosphatidic acid,^8^ have been reported to elicit its nuclear translocation. Previous reports have implied the formation of a nuclear eNOS signalosome as a complex between estrogen receptor-α and eNOS in cells treated with 17β-estradiol that associated with a specific telomerase promoter site determining chromatin remodeling and active gene transcription.^9^ Despite the wealth of work on epigenetic regulation of gene expression by NO-dependent mechanisms, what eNOS does in the nucleus has not been studied in detail. Identifying the pathways influenced by nuclear eNOS could go a long way to explaining the large influence NO has on vascular physiology and pathophysiology and eventually identify novel exploitable pathways for therapy and prevention. Therefore, this study was designed to identify nuclear proteins interacting with nuclear eNOS and elucidate the pathways regulated as a consequence.

## Methods

### Cell culture

#### Human endothelial cells

Human umbilical veins were obtained from local hospitals and endothelial cells were isolated and cultured as described previously.^10^ Experiments were performed using confluent cells that had been starved of serum overnight in endothelial cell basal medium 2 (EBM2, Cat. # C-22211; PromoCell, Heidelberg, Germany) containing bovine serum albumin (BSA) (0.1% w/v, Cat. # A8412; Sigma-Aldrich, Darmstadt, Germany), penicillin (50 units/mL), streptomycin (50 µg/mL) and sepiapterin (10 µmol/L, Cat. # S258913; LGC, Kulmbach, Germany) to maintain eNOS activity. Cells were then stimulated with human VEGF_165_ (50 ng/mL Cat. # 100-20; PeproTech, Hamburg, Germany) for 15 minutes. The use of human endothelial cells was approved by Ethic-Commission of Goethe University Frankfurt and conforms to the principles outlined in the Declaration of Helsinki.^11^

#### Murine lung endothelial cells

Pulmonary endothelial cells were isolated from wild-type mice and cultured as described.^12^ Before stimulation with murine VEGF_165_ (30 ng/mL, Cat. #450-32; PeproTech, Hamburg, Germany) for 15 minutes, cells were starved of serum overnight in in a medium containing Ham’s F-12 Nutrient Mix (50% v/v), DMEM (50% v/v), L-glutamine (1.5 mmol/L), BSA (0.1% w/v) and sepiapterin (10 µmol/L).

### Functional assays

#### Cell proliferation

Human endothelial cells were seeded into 96 well plates (Cat. # 83.3924; SARSTEDT, Nümbrecht, Germany) at a density of 4000 cells per well. After 2 hours, the medium was replaced with fresh ECGM2 containing 10 µmol/L sepiapterin. Four images/well were acquired every 3 hours using an Incucyte S3 live-cell imaging and analysis system (Essen BioScience, Hertfordshire, United Kingdom) with a 10x magnification for up to 3 days. The total number of cells per square millimeter was quantified from the images at different time points.

#### Cell death

Human endothelial cells were seeded into 96-well plates at a density of 35,000 cells per well. After 4-5 hours, the medium was replaced with fresh ECGM2 containing sepiapterin (10 µmol/L) and propidium iodide (10 µmol/L) (Cat. # P4170; Sigma-Aldrich, Darmstadt, Germany). Cells were then treated with either solvent, tumour necrosis factor-α (TNF-α; 1 ng/mL; Cat. # 300-01A; PeproTech, Hamburg, Germany) or hydrogen peroxide (H_2_O_2_, 1 µmol/L; Cat. # 8070; Roth, Karlsruhe, Germany). Four images/well were taken every 3 hours using Incucyte S3 live-cell imaging and analysis system with a 10x magnification for up to 3 days, and the number of propidium iodide-positive cells at different time points was quantified.

### Generation of adenoviruses and adenoviral transduction of human endothelial cells

Adenoviruses encoding C-terminal Flag-tagged wild-type human full-length eNOS (Ad.Flag-eNOS), enhanced green fluorescent protein (Ad.eGFP), as well as small interfering RNA targeting human eNOS (Ad.si-eNOS) were generated as described previously.^13,14^ Human endothelial cells (80-90% confluent) were transduced with 100 MOI of Ad.Flag-eNOS, Ad.eGFP or Ad.si-eNOS in EBM2 containing BSA (0.1% w/v), penicillin (50 units/mL) and streptomycin (50 µg/mL). After 16 hours, the medium was replaced with ECGM2 and cells were maintained in culture for further experiments.

To visualize eNOS, human umbilical vein endothelial cells transduced with Ad.Flag-eNOS were cultured in 8 well chamber slides (Cat. # 80826-90; Ibidi, Martinsried, Germany), serum starved overnight in EBM2 containing BSA (0.1% w/v) and sepiapterin (10 µmol/L), and then stimulated with solvent or human VEGF_165_ (50 ng/mL) for 15 minutes at 37°C.

To visualize dsRNA in the presence or absence of eNOS, human endothelial cells transduced with Ad.eGFP or Ad.si-eNOS and cultured in 8 well chamber slides were serum starved overnight in EBM2 containing BSA (0.1% w/v) and sepiapterin (10 µmol/L), followed by stimulation with human VEGF_165_ (50 ng/mL) for 24 hours.

To visualize MAVS in the presence or absence of eNOS, human endothelial cells transduced with Ad.eGFP or Ad.si-eNOS were cultured in 8-well ibidi slides in ECGM2 supplemented with sepiapterin (10 µmol/L) for 48 hours.

### Immunofluorescence

Cells were washed with phosphate buffered saline (PBS) containing 0.8 mmol/L CaCl_2_ and 1.4 mmol/L MgCl_2_ and fixed with 4% formaldehyde (Cat. # P087.1; Carl Roth, Karlsruhe, Germany) for 15 minutes at room temperature. Permeabilization and blocking was performed with a buffer containing 0.1% Triton X-100 (Cat. # 3051.2; Carl Roth, Karlsruhe, Germany) and 5% donkey serum in PBS for 30 minutes at room temperature. Cells were incubated with primary antibodies against FLAG (2 µg/mL, Cat. # F3165; Sigma-Aldrich, Darmstadt, Germany), dsRNA (2.5 µg/mL, clone J2, Cat. # 10010200; Nordic-MUbio, Eching, Germany), or MAVS (0.07 µg/mL, Cat. # 24930; Cell Signaling Technology, Leiden, Netherlands) in PBS containing 0.01% Triton X-100 and 0.5% (w/v) donkey serum for 2 hours at room temperature. Cells were then incubated with donkey anti-mouse Alexa Fluor 488 (6.67 µg/mL, Cat. # A-21202; Invitrogen, Darmstadt, Germany), donkey anti-mouse Alexa Fluor 647 (5 µg/mL, Cat. # A-31571; Invitrogen, Darmstadt, Germany), or donkey anti-rabbit Alexa Fluor 555 (6.67 µg/mL, Cat. # A-32794; Invitrogen, Darmstadt, Germany) and 4′,6-diamidino-2-phenylindole (DAPI) (0.1 µg/mL, Cat. # A1001; AppliChem, Darmstadt, Germany) diluted in PBS for 1 hour at room temperature. Finally, cells were overlaid with a mounting medium containing 50% (v/v) glycerol (Cat. # 3783.1; Carl Roth, Karlsruhe, Germany) and dithiothreitol (DTT, 1 mol/L) (Cat. # A2948; AppliChem, Darmstadt, Germany) in PBS. Images were acquired using a confocal microscope (LSM-780; Zeiss, Jena, Germany) and ZEN software (Zeiss). Image processing was done with ImageJ.

#### *En face* staining of mouse aorta

Mice were perfusion-fixed with 4% formaldehyde through the left ventricle. Connective tissue surrounding the aorta was carefully removed under a microscope (Zeiss SteREO Discovery.V8; Carl Zeiss, Oberkochen, Germany). The aorta was permeabilized in Triton X-100 (0.01%) in PBS for 1 hour, followed by blocking in donkey serum (5%) and BSA (3% w/v, Cat. # 8076.3; Carl Roth, Karlsruhe, Germany) in PBS for 1 hour at room temperature. Samples were incubated with primary antibodies against eNOS (1.25 µg/mL, Cat. # 610297; BD Biosciences, Heidelberg, Germany) and CD31 (0.08 µg/mL, Cat. # 550274; BD Biosciences, Heidelberg, Germany) in donkey serum (0.5%) and BSA (0.3% w/v) in PBS for 2 hours at room temperature. The aorta was then incubated with donkey anti-mouse Alexa Fluor 647 (10 µg/mL, Cat. # A-31571; Invitrogen, Darmstadt, Germany) and donkey anti-rat DyLight 550 (5 µg/mL, Cat. # SA5-10027; Invitrogen, Darmstadt, Germany) for 1 hour followed by incubation with Hoechst (2 µmol/L, Cat. # 94403; Sigma-Aldrich, Darmstadt, Germany) for 10 minutes at room temperature. Aorta segments were cut and placed on a microscope slide (Cat. # 0810401; Paul Marienfeld, Lauda-Königshofen, Germany) with endothelium layer exposed. Fluoromount-G mounting medium (Cat. # 00-4958-02; Thermo Fisher Scientific, Darmstadt, Germany) was added on top of each segment, before covering with a coverslip (Cat. # 15737592; Thermo Fisher Scientific, Darmstadt, Germany). Images were taken using a confocal microscope (Stellaris 5; Leica, Mannheim, Germany) and LAS X software (Leica). Image processing was done with Imaris 10.

#### Murine carotid arteries and human carotid atherosclerotic plaques

Murine carotid arteries were removed 2 days after partial ligation, cleaned from connective tissue, embedded in Tissue-Tek OCT compound (Cat. # 25608-930; VWR, Darmstadt, Germany) and frozen at - 80 °C until used. Cryosections (5-10 μm) were prepared and washed twice with PBS, before being fixed in formaldehyde (4%, 10 minutes). Formalin-fixed, paraffin-embedded sections of human carotid arteries were obtained from the Munich Vascular Biobank (Klinikum rechts der Isar, Technical University Munich) with local Ethics Committee approval (# 2799/10) and in accordance with the Declaration of Helsinki. The characteristics of patients whose carotid sections were used are available in **Supplementary Table S1**. Samples were incubated at 60°C for 10 minutes followed by deparaffinization in 100% xylene, transferred to a series of descending ethanol concentrations (100%, 96%, 90%, 80%, 70% and 50%; 5 minutes each) and finally to distilled water. Thereafter, samples were incubated with pre-heated antigen retrieval solution (10 mmol/L sodium citrate pH 6) at 90-100°C for 20 minutes. Sections were incubated with blocking buffer containing donkey serum (5%), BSA (1%), Triton X-100 (0.1%) in PBS for 30 minutes. Samples were then incubated with primary antibodies in blocking buffer overnight at 4°C, followed by secondary antibodies and DAPI (0.1 µg/mL) in PBS for 2 hours. Sections were mounted with Fluoromount-G and covered with the coverslip. The following primary antibodies were used; dsRNA (5 µg/mL, clone 9D5, Cat. # Ab00458-23.0; Absolute Antibody, Eching, Germany), OAS3 (9 µg/mL, Cat. # 21915-1-AP; Proteintech, Planegg-Martinsried, Germany), CD31 (0.078125 µg/mL, Cat. # 550274; BD Biosciences, Heidelberg, Germany) and biotinylated Ulex Europaeus Agglutinin I (10 µg/mL, Cat. # B-1065-2; Vector Laboratories, Eching, Germany). Secondary antibodies were donkey anti-rabbit Alexa Fluor 555 (6.67 µg/mL, Cat. # A-32794; Invitrogen, Darmstadt, Germany), donkey anti-rat Alexa Fluor 647 (6.67 µg/mL, Cat. # A-78947; Invitrogen, Darmstadt, Germany) or Alexa Flour 647-conjugated streptavidin (6.67 µg/mL, Cat. # S32357; Thermo Fisher Scientific, Darmstadt, Germany).

Z-stack images of mouse carotid arteries were taken using a confocal microscope (SP8; Leica, Wetzlar, Germany) and LAS AF lite software (Leica). Images of human carotid arteries were taken using a confocal microscope (LSM-780; Zeiss, Jena, Germany) and ZEN software (Zeiss). Image processing and analysis were done with ImageJ. The CD31 positive endothelium was used to create a selection for quantification of the mean gray values of the endothelial dsRNA or OAS3 signal in the thresholded image of the dsRNA or OAS3 channels, respectively.

### Protein isolation for immunoblotting

Human endothelial cells were lysed in buffer containing Tris/HCl pH 7.5 (50 mmol/L), NaCl (150 mmol/L), NaPPi (10 mmol/L), NaF (20 mmol/L), sodium deoxycholate (1% w/v), Triton X-100 (1% v/v), sodium dodecyl sulfate (SDS, 0.1% w/v) and protease and phosphatase inhibitors (okadaic acid (10 nmol/L), Na_3_VO_4_ (2 mmol/L), antipain (2 mg/L), aprotinin (2 mg/L), chymostatin (2 mg/L), leupeptin (2 mg/L), pepstatin (2 mg/L), trypsin inhibitor (2 mg/L), and phenylmethylsulfonyl fluoride (230 µmol/L)). Cell lysates were sonicated on ice (1 second on, second off pulse, 50 % amplitude, 10 cycles in total) using an Ultrasonic Homogenizer mini20 (BANDELIN, Berlin, Germany). After centrifugation (16,000 g, 5 minutes), the supernatants were recovered.

### Nuclear fractionation

For the separation of cytosol and nuclear fractions, cells were lysed in a buffer containing HEPES pH 7.6 (10 mmol/L), KCl (10 mmol/L), CaCl_2_ (1 mmol/L), NaPPi (10 mmol/L), NaF (20 mmol/L), β-glycerolphosphate (50 mmol/L), okadaic acid (1 µmol/L), Na_3_VO_4_ (2 mmol/L), phenylmethylsulfonyl fluoride (230 µmol/L), glycerol (10% v/v), Nonidet P-40 (NP-40, 0.1%) and protease inhibitors (10 minutes, 4°C). After centrifugation (10,000 g, 5 minutes), supernatants (cytosolic fractions) were transferred to fresh tubes, and the cell pellet was washed 3 times with lysis buffer (without NP-40) and centrifuged (2,500 g, 5 minutes, 4°C). Pellets were then resuspended in a buffer containing HEPES pH 7.6 (20 mmol/L), NaCl (150 mmol/L), MgCl_2_ (1.5 mmol/L), CaCl_2_ (1 mmol/L), NaPPi (10 µmol/L), NaF (20 mmol/L), β-glycerolphosphate (50 mmol/L), okadaic acid (1 µmol/L), Na_3_VO_4_ (2 mmol/L), phenylmethylsulfonyl fluoride (230 µmol/L), glycerol (10% v/v) and protease inhibitors and incubated for 10 minutes on ice with vortexing, followed by sonication as above. After centrifugation (16,000 g, 5 minutes), the supernatants (nuclear fractions) were recovered and frozen until analyzed further.

### Immunoprecipitation

Nuclear lysates (100-300 µg) were incubated with mouse IgG (0.2-0.6 µg, Cat. # sc-2025; Santa Cruz Biotechnology, Heidelberg, Germany), or antibodies directed against ADAR1 (Cat. # sc-73408; Santa Cruz Biotechnology, Heidelberg, Germany), eNOS (Cat. # 610297; BD Biosciences, Heidelberg, Germany) or FLAG (M2 Affinity Gel, Cat. # A2220; Merck, Darmstadt, Germany) overnight (4°C). Thereafter, protein G-sepharose 4B conjugate (25 µL unpacked protein G per 1 mg lysate; Cat. # 17061801; Cytiva, Dreieich, Germany) was incubated overnight (4°C) on an end-over-end rocker. After centrifugation (2,300 g, 4°C, 2 minutes), immunoprecipitates were washed 3 times with ice-cold lysis buffer and SDS-PAGE sample buffer (35 µL) was added and heated at 95°C for 5 minutes. The eluted immunoprecipitates and the inputs (30 µg) were analyzed by immunoblotting. FLAG-tagged eNOS immunoprecipitates were washed 3 times with a buffer containing Tris/HCl pH 7.5 (50 mmol/L), NaCl (150 mmol/L) and CaCl_2_ (1 mmol/L) before processing for LC-MS/MS.

### Identification of the nuclear eNOS interactome by LC-MS/MS

FLAG-tagged eNOS immunoprecipitates were supplemented with guanidinium chloride (6 mol/mL), Tris/HCl (50 mmol/L) and Tris(2-carboxyethyl)phosphine hydrochloride (TCEP, 10 mmol/L) and incubated at 95°C for 5 minutes. Reduced thiols were alkylated with chloroacetamide (40 mmol/L) and sample were diluted with 25 mmol/L Tris/HCl, pH 8.5, 10% acetonitrile to obtain a final guanidinium chloride concentration of 0.6 mol/L. Proteins were digested with 1 µg trypsin (sequencing grade, Promega) overnight at 37°C under gentle agitation. Digestion was stopped by adding trifluoroacetic acid to a final concentration of 0.5 %. Peptides were loaded on StageTip containing six C18-disks. Purification and elution of peptides was performed as described in Rappsilber and Mann (2007). Peptides were eluted in wells of microtiter plates and peptides were dried and resolved in 1% acetonitrile, 0.1 % formic acid. Liquid chromatography / mass spectrometry (LC/MS) was performed on Thermo Scientific Q Exactive Plus equipped with an ultra-high performance liquid chromatography unit (Thermo Scientific Dionex Ultimate 3000) and a Nanospray Flex Ion-Source (Thermo Scientific). Peptides were loaded on a C18 reversed-phase precolumn (Thermo Scientific) followed by separation on a with 2.4 µm Reprosil C18 resin (Dr. Maisch GmbH) in-house packed picotip emitter tip (diameter 100 µm, 15 cm from New Objectives) using a gradient from 4% acetonitrile, 0.1% formic acid to 40 % eluent B (99% acetonitrile, 0.1% formic acid) for 30 min followed by a second gradient to 60%B for 5 min with a flow rate 400 nL/minute. MS data were recorded by data dependent acquisition. The full MS scan range was 300 to 2000 m/z with resolution of 70000, and an automatic gain control (AGC) value of 3*10E6 total ion counts with a maximal ion injection time of 160 ms. Only higher charged ions (2+) were selected for MS/MS scans with a resolution of 17500, an isolation window of 2 m/z and an automatic gain control value set to 10E5 ions with a maximal ion injection time of 150 ms. MS1 Data were acquired in profile mode. MS Data were analyzed by MaxQuant (v1.6.1.0) [19029910] using default settings. Proteins were identified using reviewed human reference proteome database UniProtKB with 71785 entries, released in 2/2018. The enzyme specificity was set to Trypsin. Acetylation (+42.01) at N-terminus and oxidation of methionine (+15.99) were selected as variable modifications and carbamidomethylation (+57.02) as fixed modification on cysteines. False discovery rate (FDR) for the identification protein and peptides was 1%. Label free quantification (LFQ) and intensity-based absolute quantification (IBAQ) values were recorded. Data were further analyzed by Perseus (v. 1.6.1.3). Contaminants and reversed identification were removed. Protein identification with at least 4 valid quantification values in at least one group were further analyzed. Missing values were replaced with the lowest quantification values of the dataset. Student’s t-test was used to identify significant enriched proteins between experimental groups. Significant protein candidates supported by weak evidence defined as at least two of the following conditions: 1) one unique peptide; 2) sequence coverage lower than 3%; 3) molecular weight higher than 35 kDa were excluded from the final list of eNOS interacting proteins.

### Immunoblotting

Protein concentration of cell lysates was determined by Bradford assay. Protein samples were denatured by heating at 95°C for 5 minutes in 1X SDS-PAGE sample buffer containing glycerol (8.5% v/v), SDS (2% w/v), Tris/HCl pH 6.8 (62.7 mmol/L), DTT (20 mmol/L), and bromophenol blue (0.002 % w/v) diluted in ddH_2_O. Protein samples were then separated in SDS-PAGE gels, and transferred to 0.45 mm nitrocellulose membranes (GE Healthcare, Freiburg, Germany). Membranes were blocked with 1X ROTI Block (Cat. # A151.3; Carl Roth, Karlsruhe, Germany) diluted in ddH_2_O for 2 hours at room temperature and were incubated with primary antibodies at 4°C overnight. Primary antibodies: eNOS (0.25 µg/mL, Cat. # 610297; BD Biosciences, Heidelberg, Germany), TERT (1:1000, Cat. # 582005; Sigma-Aldrich, Darmstadt, Germany), HSP90 (0.25 µg/mL, Cat. # 610419; BD Biosciences, Heidelberg, Germany), GAPDH (1 µg/mL, Cat. # MAB374; Merck Millipore, Darmstadt, Germany), ADAR1 (0.2 µg/mL, Cat. # sc-73408; Santa Cruz Biotechnology, Heidelberg, Germany), MAVS (0.014 µg/mL, Cat. # 24930S; Cell Signaling Technology, Leiden, Netherlands), β-Actin (2.1 µg/mL, Cat. # A1978; Sigma-Aldrich, Darmstadt, Germany), OAS3 (0.027 µg/mL, Cat. # 41440; Cell Signaling Technology, Leiden, Netherlands), and RNase L (0.2 µg/mL, Cat. # sc-74405; Santa Cruz Biotechnology, Heidelberg, Germany). Thereafter, membranes were incubated with species-specific HRP-conjugated secondary antibodies for 2 hours at room temperature. Target proteins were visualized by enhanced chemiluminescence using a commercially available kit (GE Healthcare, Freiburg, Germany).

### Biotin switch technique

The biotin switch technique was performed as described,^15^ with modifications. Briefly, cells were lysed in buffer (pH 7.5) containing Tris/HCl (50 mmol/L), ethylenediaminetetraacetic acid (EDTA, 1 mmol/L), NaCl (150 mmol/L), Triton X-100 (0.4% v/v), deferoxamine (0.1 mmol/L) and neocuproine (0.1 mmol/L). All of the following steps were performed in the dark. Cell lysates were sonicated (1 second ON + 1 second OFF pulse, 50% amplitude, 15 cycles in total) and proteins precipitated in acetone (-80°C, 1 hour) and recovered by centrifugation (2,000 g, 4°C, 20 minutes). Protein pellets were subsequently dissolved in a buffer (pH 7.7) containing Tris/HCl (20 mmol/L), SDS (2.5% w/v), EDTA (1 mmol/L) and neocuproine (0.1 mmol/L). Following sonication, samples were incubated with methyl methanethiosulfonate (MMTS, 1 mol/L, Cat. # 23011; Thermo Fisher Scientific, Dreieich, Germany) at 50°C for 30 minutes to block free thiols, followed by two precipitations in acetone to remove MMTS. Pellets were then resuspended in buffer (pH 7.6) containing Tris/HCl (20 mmol/L), sonicated and split into 2 equal parts. One part was supplemented with sodium ascorbate (200 µmol/L, Cat. # 05097; Merck, Darmstadt, Germany) and HPDP-Biotin (5 µmol/L, Cat. # 21341; Thermo Fisher Scientific, Darmstadt, Germany). The second served as negative control and was treated with DTT (0.1 mol/L, Cat. # A2498; Applichem, Darmstadt, Germany), TCEP (0.1 mol/L, Cat. # C4706, Sigma-Aldrich, Darmstadt, Germany) and HPDP-Biotin (5 µmol/L). Samples were then briefly vortexed and incubated for 1 hour at room temperature under gentle agitation before proteins were precipitated with acetone. Precipitated proteins were suspended in incubation buffer (pH 7.7) containing Tris/HCl (20 mmol/L), EDTA (1 mmol/L), and Triton X-100 (0.4% v/v) with sonication. Some protein (30 µg) was retained as the “input”, and 200-300 µg protein was immunoprecipitated by incubated with streptavidin agarose (Cat. # 20359; Thermo Fisher Scientific, Darmstadt, Germany) under constant agitation (room temperature, 2 hours) and centrifugation (1,000 g, 10 minutes). Biotinylated proteins were eluted by incubation with 2x SDS-PAGE sample buffer containing β-mercaptoethanol (100 µmol/L, Cat. # M6250; Sigma-Aldrich, Darmstadt, Germany) for 20 minutes at room temperature, followed by heating at 95°C for 10 minutes. The eluted samples and the inputs were analyzed by immunoblotting.

### RNA isolation

To knockdown eNOS in human endothelial cells, Ad.si-eNOS (100 MOI) was used to transduce cells grown in 6-well plate (eNOS KD). Endothelial cells transduced with Ad.eGFP served as control (CTL). Thirty-six hours after transduction, the cells were serum starved in EBM2 containing sepiapterin (10 µmol/L) for 8 hours, followed by stimulation with human VEGF_165_ (50 ng/mL) for additional 6 hours. Cells were then lysed in lysis buffer containing Tris/HCl pH 7.5 (50 µmol/L), NaCl (150 µmol/L), MgCl_2_ (12 µmol/L), NP-40 (1% v/v), NaPPi (10 mmol/L), NaF (20 µmol/L), β-glycerolphosphate (50 µmol/L), protease and phosphatase inhibitors, SUPERase-In RNase Inhibitor (200 U/mL, Cat. # AM2696; Thermo Fisher Scientific, Darmstadt, Germany), and TURBO DNase (25 U/mL, Cat. # AM2239; Thermo Fisher Scientific, Darmstadt, Germany). Lysates were collected into RNase-free tubes and incubated on ice for 10 minutes. Homogenization was performed by passing cell lysates several times through a 26G needle, and the supernatants were recovered by centrifugation (13,000 g, 4°C, 10 minutes). The supernatants were transferred into fresh RNase-free tubes and total RNA was purified using the miRNeasy Micro kit (Cat. # 217684, QIAGEN, Hilden, Germany), according to the manufacturer’s instructions.

### Next generation sequencing

Total RNA (1 µg) was used as input for SMARTer Stranded Total RNA Sample Prep Kit - HI Mammalian (Cat. # 634875; Takara, Saint-Germain-en-Laye, France). Sequencing was performed on the NextSeq 2000 instrument (Illumina) using P3 flow cell and 1x72 bp single end setup. Trimmomatic version 0.39 was employed to trim reads after a quality drop below a mean of Q15 in a window of 5 nucleotides and keeping only filtered reads longer than 15 nucleotides.^16^ Reads were aligned versus Ensembl human genome version hg38 (Ensembl release 109) with STAR 2.7.10a.^17^ Alignments were filtered to remove: duplicates with Picard 3.0.0 (Picard: A set of tools (in Java) for working with next generation sequencing data in the BAM format), multi-mapping, ribosomal, or mitochondrial reads. Gene counts were established with featureCounts 2.0.4 by aggregating reads overlapping exons on the correct strand excluding those overlapping multiple genes.^18^ The raw count matrix was normalized with DESeq2 version 1.36.0.^19^ Contrasts were created with DESeq2 based on the raw count matrix. Genes were classified as significantly differentially expressed at average count > 5, multiple testing adjusted P value < 0.05, and log2 fold change > 0.585. The Ensemble annotation was enriched with UniProt data. Differentially expressed transcripts (FDR ≤ 0.05) following eNOS knockdown were used for gene set enrichment analysis (Kobas,^20^ Reactome, FDR ≤ 0.05).

### Adenosine-to-inosine (A-to-I) editing analysis

Trimmed, mapped and filtered reads of the RNA-Seq analysis were used to explore differential A-to-I editing by comparing conditions with JACUSA version 2.0.1 parameters “call-2 -a D,M,Y-filterNM 5 -s -c 2 -T 1.56 -P FR-SECONDSTRAND”.^21^ Events were filtered to only keep reads with < 5 mismatches and editing sites with > 2 supporting reads, and Jacusa likelihood score > 1.56. Editing sites with a modification ratio > 0.95 were removed to exclude putative genomic single nucleotide polymorphisms. The relative A-to-I editing ratio for each site in RNA transcripts in one group referred to its transformed ratio normalized to that in the other group, and the transformed ratio = (the number of G reads + 1) / (the number of A reads + 1 + the number of G reads + 1). The number of more frequently A-to-I edited sites of each transcript (log2 editing ratio > 0) was compared to its expression level in the presence/absence of eNOS in order to explore the correlation between editing and gene expression (FDR ≤ 0.05).

### Enzyme-linked immunosorbent assay (ELISA)

Endothelial cell supernatant was collected and centrifuged (2,000 g, 10 minutes) to remove any cell debris before being transferred to fresh tubes and freeze-dried overnight. The lyophilized products were dissolved in 50-150 µL sample diluent and the levels of IFN-α or IFN-β were measured with ELISA kits (IFN-α: Cat. # ab213479; Abcam, Amsterdam, Netherlands; IFN-β: Cat. # ab278127; Abcam, Amsterdam, Netherlands) according to the manufacturer’s instructions.

### siRNA transfection of human endothelial cells

To silence ADAR1, human endothelial cells were transfected with small interfering RNA (siRNA) directed against ADAR1 (5′ GCU-AUU-UGC-UGU-CGU-GUC-AdTdT 3′ and 5′ UGA-CAC-GAC-AGC-AAA-UAG-CdTdT 3′; Eurogentec, Seraing, Belgium), or with control oligonucleotides (5′ UUC-UCC-GAA-CGU-GGC-ACG-AdTdT 3′ and 5′ UCG-UGC-CAC-GUU-CGG-AGA-AdTdT 3′; Eurogentec, Seraing, Belgium). When endothelial cells reached 80% confluency, control siRNA or siRNA targeting ADAR1 (30 pmol per well in 6-well plate) and Lipofectamine RNAiMAX transfection reagent (9 µL; Cat. # 13778075; Thermo Fisher Scientific, Dreieich, Germany) were used to transfect the cells according to manufacturer’s instructions. After 4-6 hours, the medium was replaced with fresh ECGM2 and experiments performed after a further 48 hours.

### Multiplexed enhanced protein dynamic mass spectrometry

Human endothelial cells expressing and lacking eNOS and ADAR1 were starved of serum in EBM2 containing BSA (0.1% w/v) and sepiapterin (10 µmol/L) for 16 hours before the addition of heavy labelled amino acid DMEM medium (SILAC) (Cat. # 88364; Thermo Fisher Scientific, Rockford, USA) containing 5% FBS and sepiapterin (10 µmol/L) for a further 6 hours. Thereafter, cells were then collected by trypsinisation, washed with PBS and lysed as described previously.^22^ Cells were lysed and denatured with SDS (2% w/v), Tris/HCl pH 8 (50 µmol/L), NaCl (150 µmol/L), TCEP (10 µmol/L), chloroacetamide (40 µmol/L) and EDTA-free protease inhibitor cocktail (Cat. # 11836170001; Roche, Mannheim, Germany) at 95°C for 10 minutes, followed by sonication for 1 minute (1 second ON + 1 second OFF pulse, 45% amplitude, 30 cycles) and second heating at 95°C for 5 minutes. Pure proteins were obtained with methanol/chloroform precipitation. Protein pellets were then resuspended in urea. Protein (50 μg per sample) was digested overnight at 37°C with LysC (Wako Chemicals) at 1:50 (w/w) ratio and trypsin (Promega, V5113) at 1:100 (w/w) ratio. Digested peptides were purified using Sep-Pak tC18 cartridges (Waters, WAT054955). Peptide concentrations were determined with a μBCA assay (Cat. # 23235; Thermo Fisher Scientific, Darmstadt, Germany) and 10 μg of peptide per sample was labeled with TMTpro18 plex reagents (Cat. # A52045; Thermo Fisher Scientific, Darmstadt, Germany). TMT-labeled samples were adjusted to equal amounts and pooled. Then the pool was fractionated into 8 fractions using the High pH fractionation Thermo-Kit (Cat. # 84868; Thermo Fisher Scientific, Darmstadt, Germany) method. For High pH fractionation, High pH Reversed-Phase Peptide columns were washed twice with 100% acetonitrile, followed by equilibration with 0.1% TFA. Peptides were loaded in 3% acetonitrile, 0.1% TFA and washed with 3% acetonitrile concentrations in 0.1 % triethylamine solution. Elution was performed with step-wise increasing acetonitrile concentrations in 0.1% triethylamine (7.5%, 10%, 12.5%, 15%, 17.5%, 20%, 22.5%, and 50%). The eight fractions were then dried by vacuum centrifugation and resuspended in 2% ACN and 0.1% TFA for LC-MS analysis. Samples were shot with settings described earlier.^23^ Briefly, fractions were resuspended in 2% acetonitrile and 0.1% formic acid and separated on an Easy nLC 1200 (Thermo Fisher Scientific) and a 35 cm long, 75 μm ID fused-silica column, which had been packed in house with 1.9 μm C18 particles (ReproSil-Pur, Dr. Maisch) and kept at 50°C using an integrated column oven (Sonation). HPLC solvents consisted of 0.1% formic acid in water (Buffer A) and 0.1% formic acid, 80% acetonitrile in water (Buffer B). Assuming equal amounts in each fraction, 1 µg of peptides were eluted by a non-linear adapted for each fraction (F1: 3-24, F2: 4-26, F3: 5-28, F4: 6-30, F5: 7-32, F6: 8-34, F7: 9-36, F8: 10-40 %B) over 150 minutes, followed by an increase to 60% B in 5 minutes then 1 minute to 95%B and held for another 9 minutes. After that, peptides directly sprayed into an Orbitrap Ascend Tribrid mass spectrometer (Thermo Fisher Scientific). A synchronous precursor selection (SPS) multi-notch MS3 method was used in order to minimize ratio compression as previously described^24^. Full scan MS spectra (350-1400 m/z) were acquired with a resolution of 120,000 at m/z 200, maximum injection time of 100 ms and AGC target value of 4x10^5^. The most intense precursors with a charge state between 2 and 5 per full scan were selected for fragmentation (“Top Speed” with a cycle time of 1.5 seconds) and isolated with a quadrupole isolation window of 0.7 Th. MS2 scans were performed in the Ion trap (Turbo) using a maximum injection time of 35 ms, AGC target value of 1x10^4^ and fragmented using CID with a normalized collision energy (NCE) of 35%. SPS-MS3 scans for quantification were performed on the 10 most intense MS2 fragment ions with an isolation window of 0.7 Th (MS) and 2 m/z (MS2). Ions were fragmented using HCD with an NCE of 65% (TMTclassic)/ 50% (TMTpro) and analyzed in the Orbitrap with a resolution of 45,000 at m/z 200, scan range of 100-200 m/z, AGC target value of 1x10^5^ and a maximum injection time of 91 ms. Repeated sequencing of already acquired precursors was limited by setting a dynamic exclusion of 60 seconds and 7 ppm and advanced peak determination was deactivated. All spectra were acquired in centroid mode. Raw data was analyzed with Proteome Discoverer 2.4 (Thermo Fisher Scientific). The SequenceHT node was selected for database searches of MS3-spectra. Human trypsin digested proteome (Homo sapiens SwissProt database (TaxID:9606, version 2023-02-14)) was used for protein identifications. Contaminants (MaxQuant “contamination.fasta”) were determined for quality control. TMTpro (+304.207) at the N-terminus, TMTproK8 (K, +312.221), Arg10 (R, +10.008) and carbamidomethyl (C, +57.021) at cysteine residues were set as fixed modifications. Methionine oxidation (M, +15.995) and acetylation (+42.011) at the protein N-terminus as well as Met-loss (M, -131.040) were set for dynamic modifications. Precursor mass tolerance was set to 20 ppm and fragment mass tolerance was set to 0.5 Da. Default Percolator settings in PD were used to filter peptide spectrum matches (PSMs). Reporter ion quantification was achieved with default settings in consensus workflow. PSMs were exported for further analysis using the DynaTMT package.^23^ Normalized abundances from DynaTMT results were used for statistical analysis after contaminations and complete empty values were removed. Significantly altered proteins were determined using the PBLMM tool.^25^ Differentially expressed proteins (FDR ≤ 0.05) following eNOS knockdown or ADAR1 knockdown were used for pathway enrichment analysis (STRING,^26^ Reactome and Gene Ontology, FDR ≤ 0.05).

### AAV-PCSK9 generation

An adeno-associated virus (AAV) serotype 8 vector encoding murine gain-of-function proprotein convertase subtilisin/kexin type 9 (PCSK9) mutant D377Y-PCSK9 cDNA (AAV-PCSK9) was produced using the two-plasmid-method by co-transfecting AAV/D377Y-PCSK9 (gift from Jacob Bentzon; Addgene plasmid, Cat. # 58376)^27^ together with the helper plasmid pDP8^28^ in HEK-293T cells using polyethylenimine (Sigma-Aldrich, Darmstadt, Germany). AAV vectors were purified using iodixanol step gradients and titrated as described.^29^

### Animals and partial carotid artery ligation

Wild-type C57BL/6 and apolipoprotein E knockout (ApoE^-/-^) mice were purchased from Charles River (Sulzfeld, Germany). Mice homozygous for the eNOS Tyr656Phe mutation (eNOS^YF^ mice) were generated and crossed back onto an ApoE^-/-^ background (ApoE-eNOS^YF^ mice) as described.^30^ Endothelial cell-specific RiboTag mice (EC-RiboTag) were generated by crossing endothelial cell-specific Cre-driver mice (Cdh5-CreERT2)^31^ with Ribo-Tag (*Rpl22*^HA/HA^)^32^ mice. The offspring were backcrossed with C57BL/6 mice for at least 8-10 generations. Cre-mediated recombination was induced in 8-9-week-old mice by intraperitoneal injections of tamoxifen (50 mg/kg per day dissolved in 50 µL Miglyol; Merck, Darmstadt, Germany) for 5 consecutive days. Three to five days after the last tamoxifen injection, EC-RiboTag mice (9-10 weeks of age) were injected once with AAV-PCSK9 (10^11^ VG) via the tail vein. All mice were fed a cholesterol rich diet (metabolizable energy 35 kJ% fat, containing 12.5 mg/kg cholesterol; EF PAIGEN, Ssniff, Soest, Germany). One week after initiation of the cholesterol rich diet, partial ligation of the left carotid artery was performed as described.^30,33^ After 2 days (ApoE^-/-^ and ApoE-eNOS^YF^ mice) or 7 days (EC-RiboTag mice), mice were sacrificed after deep anesthetization with 180 mg/kg ketamine and 16 mg/kg xylazine, and intracardiac perfusion (through the left and right ventricles) was performed with cycloheximide (10 mL 100 μg/mL; Cat. # C4859; Sigma-Aldrich, Darmstadt, Germany) in PBS (only EC-RiboTag mice). Carotid arteries were then isolated, snap frozen in liquid nitrogen and stored at -80°C until use. Animals were housed under a 12-hour light-dark cycle with free access to water and food.

All animal experiments were performed in accordance with the Directive 2010/63/EU of the European Parliament on the protection of animals used for scientific purposes and approved by the Federal Authority for Animal Research at the Regierungspräsidium Darmstadt (Hessen, Germany) under study protocols B2/1102, B2/1187, F28/42 and FU/1231. Both male and female mice were included in the studies which were performed using age-, and strain-matched animals throughout.

### RiboTag and RNA isolation

Murine carotid arteries were homogenized in lysis buffer (2-3% w/v) containing: Tris/HCl pH 7.5 (50 mmol/L), NaCl (150 mmol/L), MgCl_2_ (12 mmol/L), NP-40 (1%), DTT (1 mmol/L), EDTA-free protease inhibitor mix (AppliChem GmbH, Darmstadt, Germany), SUPERase-In RNase Inhibitor (200 U/mL, Cat. # AM2694; Thermo Fisher Scientific, Darmstadt, Germany), cycloheximide (100 µg/mL, Cat. # C4859; Sigma-Aldrich, Darmstadt, Germany), TURBO DNase (25U/mL, Cat. # AM2238; Thermo Fisher Scientific, Darmstadt, Germany), murine RNase inhibitor (500 U/mL, Cat. # M0314L; New England Biolabs, Frankfurt am Main, Germany) and heparin (10 mg/mL) in UltraPure DNase/RNase-free distilled water (Cat. # 10977035; Thermo Fisher Scientific, Darmstadt, Germany). Lysates were centrifuged (16,000 g, 10 minutes, 4°C) and the supernatants passed through 70 µm pre-separation filters (Miltenyi Biotec, Bergisch Gladbach, Germany). A fraction (10%) of the total lysate volume was mixed with 9 volumes of QIAZOL (Cat. # 79306; QIAGEN, Hilden, Germany) and frozen at -80°C. The remaining lysates were incubated with anti-HA antibody-conjugated magnetic beads (50 µL/sample, washed and resuspended in lysis buffer; Biozol, Eching, Germany) before being incubated overnight on end-over-end rocker at 4°C. HA immunoprecipitates were washed 3 times with a freshly prepared high salt buffer containing: Tris/HCl pH 7.5 (50 mmol/L), NaCl (300 mmol/L), MgCl_2_ (12 mmol/L), NP-40 (0.5%), DTT (1 mmol/L), SUPERase-In RNase Inhibitor (200 U/mL) and cycloheximide (100 µg/mL), murine RNase inhibitor (500 U/mL) was also included. After the final wash, HA immunoprecipitates were suspended in 700 µL QIAZOL and immediately processed for RNA purification. The total input RNA (from the total lysates) as well as the ribosome-associated RNA (from the RiboTag immunoprecipitates) were purified using the miRNeasy Micro kit (QIAGEN, Hilden, Germany) according to the manufacturer’s instructions. Next generation sequencing of endothelial cell-specific transcripts was performed as described above.

### Statistics

Data are presented as mean ± standard error of the mean (SEM). GraphPad Prism software (version 10, San Diego, USA) was used to analyze the data. For data that passed a Shapiro-Wilk normality test, differences between two groups were compared by two-tailed unpaired t-test. All experiments in which the effects of two variables were tested were analyzed by 2 way ANOVA followed by the Šídák’s multiple comparisons test. Data which did not pass Shapiro-Wilk normality test were processed using the Mann-Whitney test. Differences were considered statistically significant when P<0.05.

### Data and materials availability

The mass spectrometry proteomics data have been deposited to the ProteomeXchange Consortium via the PRIDE^34^ partner repository with the dataset accession PXD052482 and PXD051054. NGS data have been deposited to the Gene Expression Omnibus and the accession numbers are GSE264495 and GSE195453.

### Datasets

Dataset 1_Nuclear eNOS interactome: nuclear eNOS interactome in human endothelial cells after stimulation with VEGF or solvent.

Dataset 2_*S*-nitrosation of eNOS interacting proteins: published *S*-nitrosated proteins, detected *S*-nitrosated cysteine residues, and related references.

Dataset 3_A-to-I editome_eNOS KD: RNA transcripts with more frequently edited sites in human endothelial cells in the presence or absence of eNOS.

Dataset 4_NGS HUVEC_eNOS KD: gene expression following eNOS knockdown in human endothelial cells.

Dataset 5_Proteome_eNOS KD_ADAR1 KD: proteins detected by mass spectrometry following eNOS or ADAR1 knockdown in human endothelial cells.

Table S1_Characteristics atherosclerosis patients: information of patients whose carotid atherosclerotic plaques were stained for visualizing dsRNA and OAS3 in the endothelium.

## Results

### Nuclear eNOS interacts with RNA-binding proteins and *S*-nitrosates ADAR1

eNOS was detected in the cytosol, Golgi apparatus and nuclei of human endothelial cells and nuclear translocation was increased by cell stimulation with VEGF for 15 minutes (**Fig. 1A and 1B**). The latter observation was not a phenomenon solely associated with cell culture as eNOS was detected *ex vivo* in nuclei in isolated murine aortae (**Fig. 1C**). Next, the nuclear eNOS interactome was determined by immunoprecipitating FLAG from nuclear extracts prepared from eNOS-FLAG expressing human endothelial cells, and subjecting the immunoprecipitates to mass spectrometry. Nuclear eNOS interacted with a number of proteins under basal conditions and following stimulation with VEGF, 81 additional proteins were identified. The latter were involved in RNA binding and processing, paraspeckle and nuclear speckle formation as well as binding to double-stranded RNA (dsRNA) (**Fig. 1D and 1E**, **Supplementary dataset 1**). Sixty-four of the nuclear eNOS-interacting proteins (79%) were already known to be *S*-nitrosated in different cell types and after exposure to either endogenously generated or exogenously applied NO (**Supplementary dataset 2**). One particularly interesting member of the latter group of proteins was the dsRNA-specific adenosine deaminase (ADAR1),^35–38^ which was previously linked to cardiovascular disease.^39^ The formation of a complex between eNOS and ADAR1 was validated by co-immunoprecipitation of the two proteins from nuclear extracts of human (**Fig. 1F**) and murine endothelial cells (**Fig. 1G**). Moreover, consistent with previous reports in other cell types,^35–38^ ADAR1 was *S*-nitrosated in human endothelial cells (**Fig. 1H**).

**Fig. 1.**
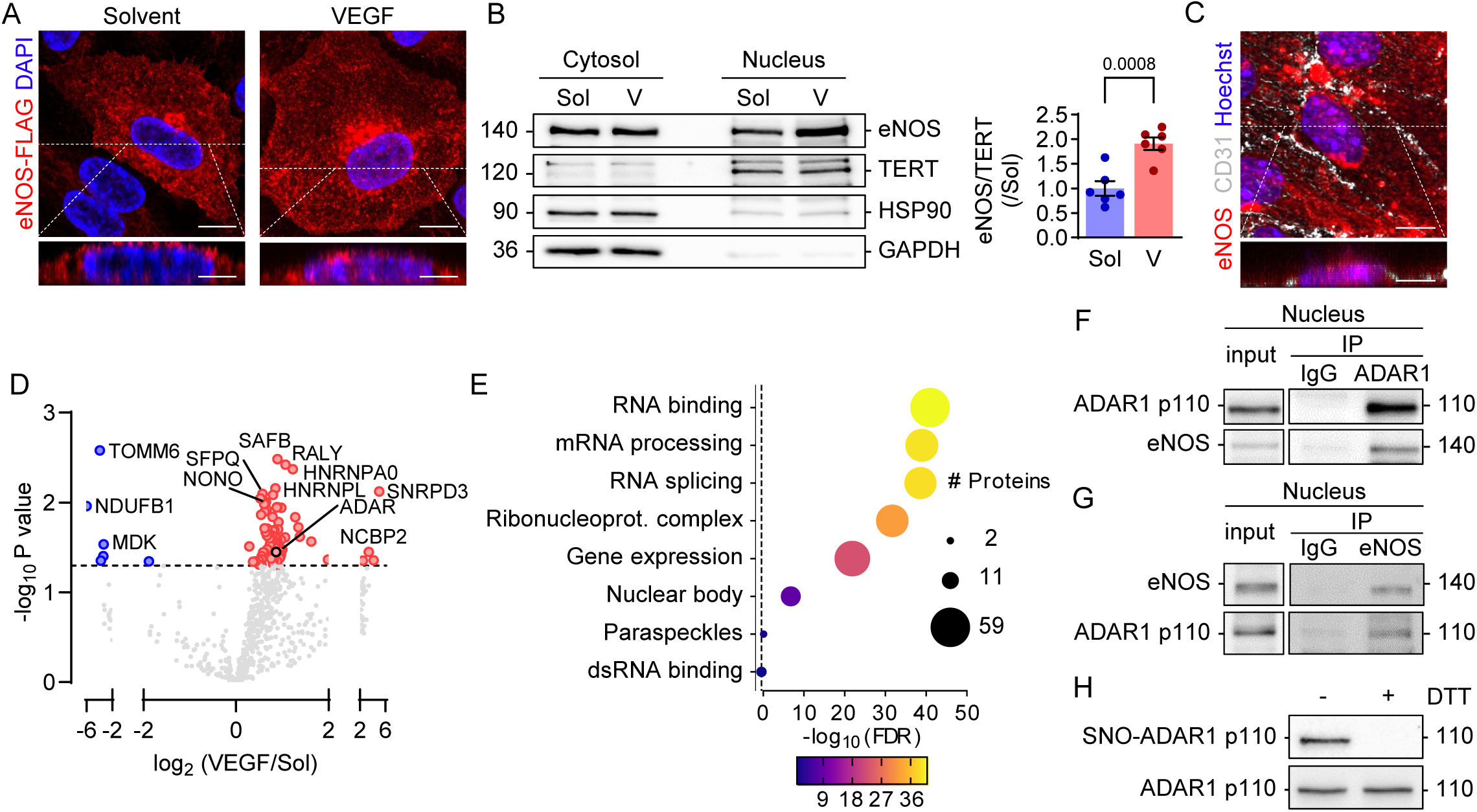
Nuclear eNOS interacts with RNA binding proteins and S-nitrosates ADAR1. (**A**) Representative confocal images of eNOS-FLAG in human endothelial cells treated with solvent (Sol) or VEGF (50 ng/ml, 15 minutes). Comparable images were obtained in 4 additional independent cell batches. Scale bars: 10 µm (xy) and 5 µm (z stack). (**B**) eNOS in the nucleus of human endothelial cells treated as in A. V=VEGF; n=6 independent cell batches (unpaired Student’s t-test). (**C**) *En face* confocal image of mouse aorta showing eNOS and CD31 *in situ*. Scale bars: 10 µm (xy) and 5 µm (z stack). (**D-E**) Volcano plot (D) and GO term enrichment analysis (STRING) (E) of the proteins physically associated with nuclear eNOS-FLAG (open circles, P value < 0.05) in human endothelial cells treated with solvent or VEGF; n=6 independent cell batches. The dashed lines mark the significance threshold (P value or FDR: 0.05). (**F**) Co-immunoprecipitation (IP) of endogenous eNOS and ADAR1 from nuclear extracts of human endothelial cells stimulated with VEGF. Similar results were obtained using 5 additional cell batches. (**G**) Co-immunoprecipitation (IP) of ADAR1 and eNOS from nuclear extracts of murine endothelial cells stimulated with VEGF (30 ng/ml, 15 minutes). Similar results were obtained using 3 additional cell batches. (**H**) *S*-nitrosation of ADAR1 in human endothelial cells. DTT was included as a negative control. Comparable results were obtained in 5 additional cell batches.

### Knockdown of eNOS results in dsRNA accumulation and alters the A-to-I editome

ADAR1 binds to dsRNA and catalyzes A-to-I editing to alter dsRNA stability.^40,41^ As the post-translational modification of ADAR1 by phosphorylation and sumoylation is known to alter its A-to-I editing function,^42,43^ we tested the hypothesis that *S*-nitrosation would exert a similar effect. Indeed, in endothelial cells the small interfering RNA-mediated depletion of eNOS significantly increased dsRNA levels (**Fig. 2A and 2B**). Moreover, when A-to-I editing frequencies were determined by next generation RNA sequencing, we found that 502 sites in 447 transcripts were more frequently edited in eNOS-deficient versus eNOS-expressing cells. In the presence of eNOS, 437 sites in 419 transcripts were more frequently edited than in its absence (**Fig. 2C**, **Supplementary dataset 3**). Edited sites were mostly located in 3′ untranslated regions (∼60%) and main coding sequences (22-24%) (**Fig. 2D**). Most transcripts presented one or two differentially edited sites and there was no obvious correlation between the frequency of editing and the expression level of the edited transcripts (**Fig. 2E**). The transcript most differentially edited by altering eNOS expression was the mitochondrial antiviral signaling protein (MAVS), which is required for the dsRNA-triggered activation of transcription factors regulating the expression of type I interferons (IFNs).^44,45^

**Fig. 2.**
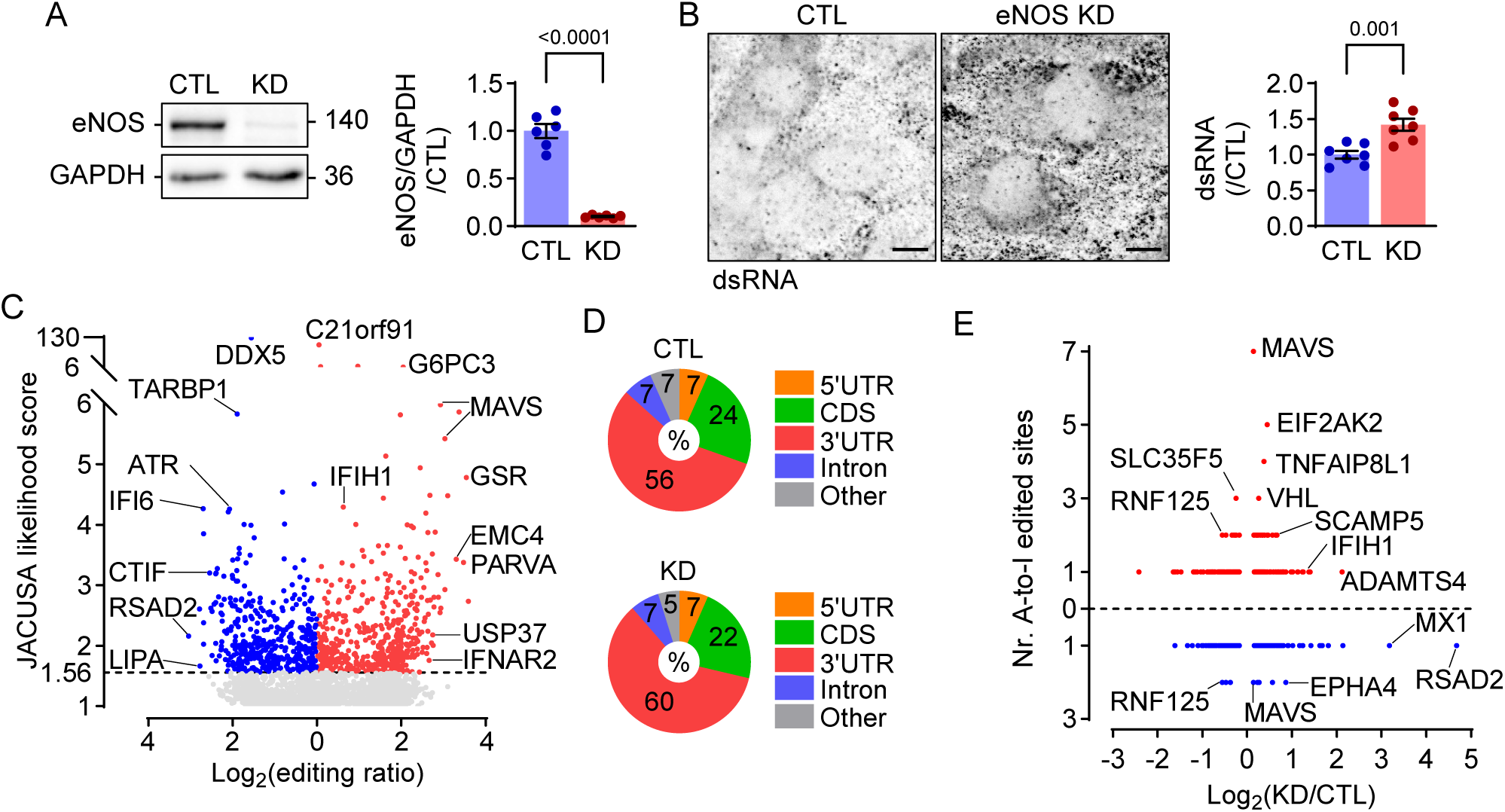
Effect of eNOS/NO signaling on the amount of dsRNA and A-to-I editing. (**A**) Efficiency of siRNA-mediated eNOS deletion (KD) in human endothelial cells; n=6 independent cell batches (unpaired Student’s t-test). **B**, Levels of dsRNA in eNOS-expressing (CTL) and eNOS-deficient (KD) human endothelial cells; scale bar: 10 µm. n=7 independent cell batches (unpaired Student’s t-test). (**C-D**) More frequently A-to-I edited sites in eNOS-expressing (blue, CTL) and eNOS-deficient (red, KD) endothelial cells (C), and their location in transcripts (D). Editing ratio values are positive as they reflect the frequency of A-to-I editing events when comparing CTL vs. KD (blue) or KD vs. CTL (red). UTR=untranslated region; CDS=coding sequence. n=4 independent cell batches. The dashed line in C marks the threshold JACUSA likelihood score: 1.56. (**E**) Dot plot depicting the relationship between the total number of A-to-I edited sites per transcript (y) and the relative expression (RNA-seq) of each transcript (x) in the presence or absence of eNOS. Red: transcripts more frequently edited in KD group, blue: transcripts more frequently edited in CTL group; n=4 independent cell batches.

### Reduced NO bioavailability triggers a dsRNA-dependent type I interferon response

MAVS is as an adaptor protein that binds to the dsRNA sensor “interferon-induced helicase C domain-containing protein 1 (IFIH1)” to mediate type I IFN responses.^46–48^ Therefore, we determined whether increasing dsRNA levels by knocking down eNOS to prevent the *S*-nitrosation of ADAR1 was able to induce a type I IFN response (**Fig. 3A**). In the absence of eNOS, the expression of MAVS as well as the formation of active MAVS clusters were consistently increased (**Fig. 3B and 3C**) as was the production of IFN-α and IFN-β (**Fig. 3D and 3E**). Gene expression analysis confirmed the significant upregulation of genes related to IFNα/β signaling as well as a marked downregulation of cell cycle-related genes in eNOS-deficient endothelial cells (**Fig. 3F and 3G**, **Supplementary dataset 4**). Indeed, proliferation was abrogated in eNOS-depleted endothelial cells (**Fig. 3H**).

**Fig. 3.**
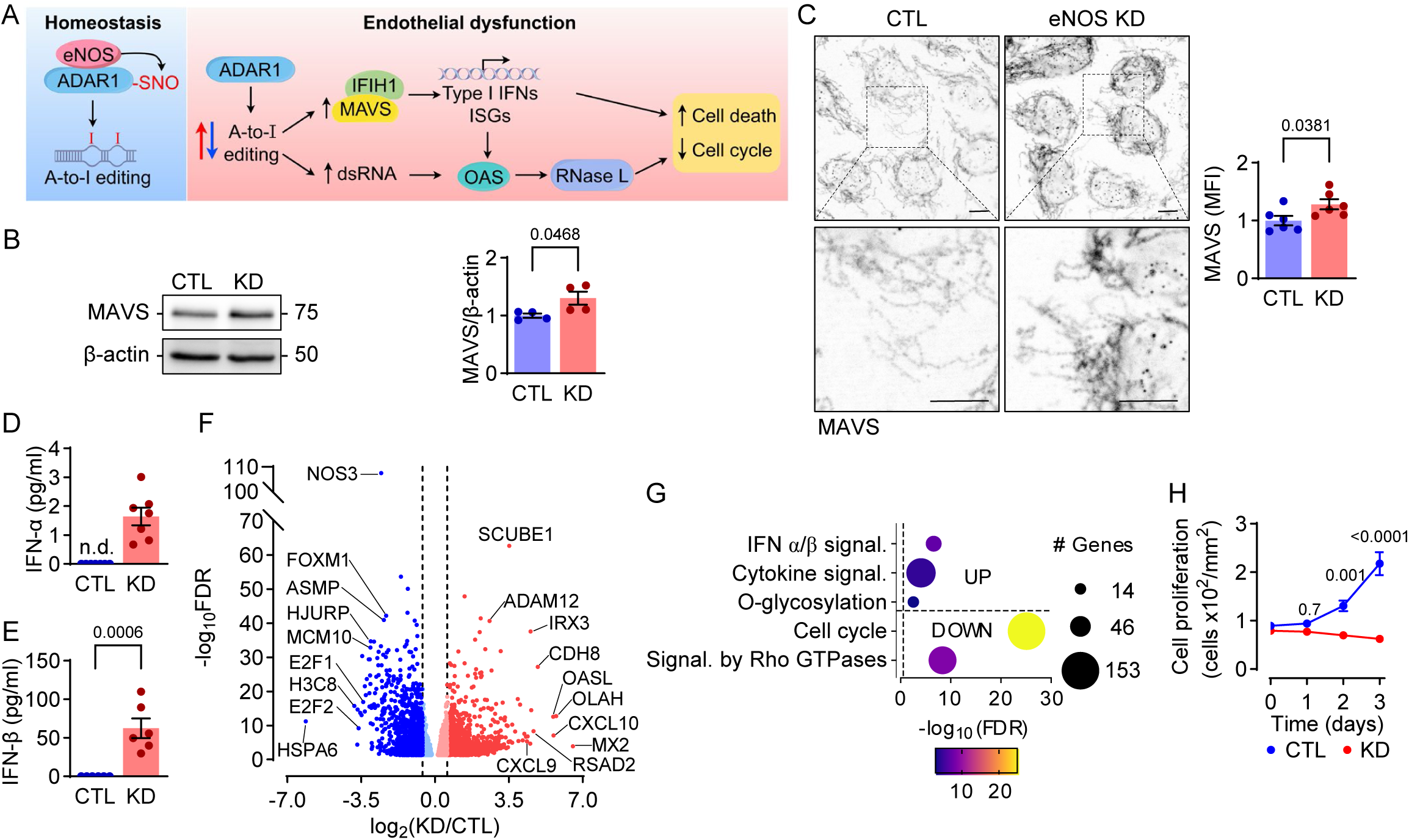
Effect of eNOS/NO signaling on the dsRNA-induced Type I IFN response. (**A**) Scheme of the proposed pathway by which a decrease in the *S*-nitrosation of ADAR1 elicits the accumulation of dsRNA to activate a type I IFN response. (**B**) Expression of MAVS in human endothelial cells in the presence (CTL) or absence (KD) of eNOS; n=4 independent cell batches (unpaired Student’s t-test). (**C**) Confocal images showing MAVS in CTL and eNOS KD endothelial cells. Scale bars: 10 µm; n=6 independent cell batches (unpaired Student’s t-test). (**D-E**) IFN-α (D) and IFN-β (E) levels in cell culture supernatants from eNOS-expressing (CTL) and –deficient (KD) endothelial cells. n.d. = not detectable; n=6-7 independent cell batches (unpaired Student’s t-test). (**F**) Volcano plot depicting the upregulated and downregulated transcripts in human endothelial cells expressing (CTL) or lacking (KD) eNOS; n=4 independent cell batches. False discovery rate (FDR) threshold: 0.05. The dashed lines mark the log_2_ fold change: ±0.585. (**G**) Gene set enrichment analysis (Kobas, Reactome) of transcripts differentially expressed following eNOS knockdown. The dashed line marks the significance threshold (FDR: 0.05). (**H**) Growth factor-induced proliferation of human endothelial cells expressing (CTL) or lacking (KD) of eNOS; n=8 independent cell batches (two-way ANOVA and Šídák’s multiple comparisons test).

The dsRNA-dependent activation of the type I IFN signaling was further assessed by mass spectrometry. This revealed that eNOS deletion significantly increased the expression of numerous dsRNA-sensitive IFN-inducible proteins. The latter included, amongst others, 2’-5’-oligoadenylate synthetase (OAS) 1 and OAS2, IFN-induced GTP-binding protein MX1, IFN-induced protein with tetratricopeptide repeats 1 (IFIT1), ubiquitin-like protein ISG15 and IFN-stimulated gene 20 kDa protein (ISG20) (**Fig. 4A**, **Supplementary dataset 5**). The increased expression of OAS3, p150 ADAR1 and p110 ADAR1 was confirmed by immunoblotting (**Fig. 4B**). The accumulation of 2’-5’-oligoadenylate generated by the OAS enzymes activates endoribonuclease RNase L to induce cell death,^49^ the expression of which was also increased in eNOS-deficient cells (**Fig. 4C**). As a consequence, eNOS knockdown increased basal as well as tumor necrosis factor-α and H_2_O_2_-induced endothelial cell death (**Fig. 4D-F**). Importantly, similar alterations in the expression of type I IFN signaling proteins were detected following the deletion of eNOS and ADAR1 (**Fig. 4G**, **Supplementary dataset 5**). Moreover, pathway enrichment analysis confirmed that type I IFN signaling was the most upregulated pathway by eNOS as well as ADAR1 deletion (**Fig. 4H and 4I**).

**Fig. 4.**
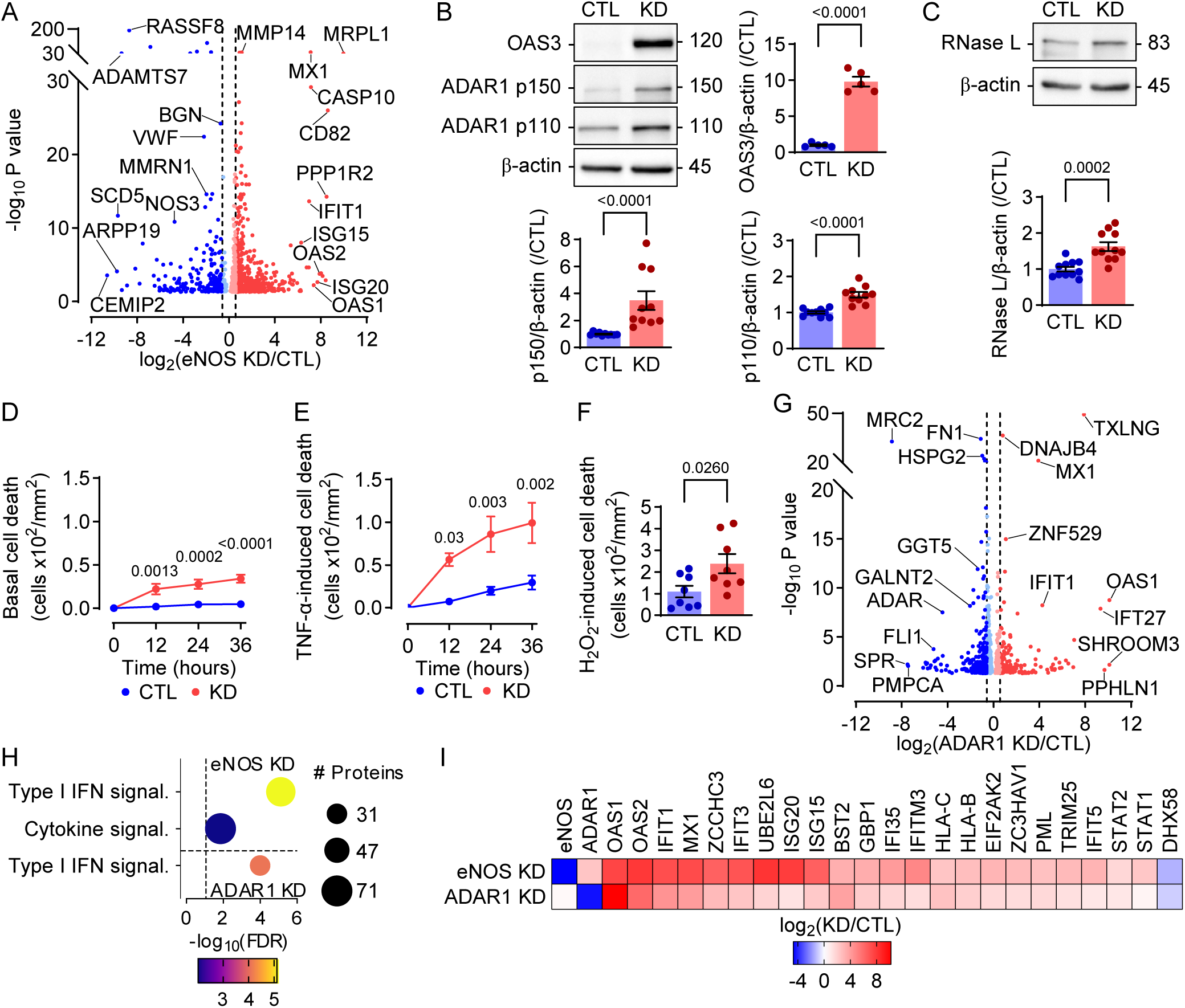
The ablation of eNOS/NO signaling phenocopies ADAR1 knockdown. (**A**) Volcano plot depicting newly synthetized proteins in human endothelial cells expressing (CTL) or lacking (eNOS KD) eNOS; n=4 independent cell batches (P value ≤ 0.05). The dashed lines mark the log_2_ fold change: ±0.585. (**B**) Expression of OAS3, ADAR1 p150 and ADAR1 p110 in human endothelial cells in the presence (CTL) or absence (KD) of eNOS; n=5-10 independent cell batches (unpaired Student’s t-test). (**C**) Expression of RNase L in human endothelial cells in the presence (CTL) or absence (KD) of eNOS; n=11 independent cell batches (unpaired Student’s t-test). (**D-F**) Impact of eNOS deletion (KD) on endothelial cell death under basal conditions (D), following stimulation with TNF-α (E), or stimulation with H_2_O_2_ (F); n=4 independent cell batches (two-way ANOVA and Šídák’s multiple comparisons test). (**G**) Volcano plot of newly synthetized proteins in human endothelial cells in the presence (CTL) or absence (ADAR1 KD) of ADAR1; n=3-4 independent cell batches (P value ≤ 0.05). The dashed lines mark the log_2_ fold change: ±0.585. (**H**) Gene set enrichment analysis (Reactome) of the differentially expressed proteins following knockdown of eNOS or ADAR1. The dashed line marks the significance threshold (FDR: 0.05). (**I**) Heatmap highlighting differentially expressed type I IFN signaling proteins in the datasets shown in A and G.

### Endothelial dysfunction induces type I IFN signaling in a mouse model of atherogenesis and in human atherosclerosis

Next, we assessed whether endothelial dysfunction *in vivo* was also associated with altered dsRNA levels and the activation of type I IFN signaling. To detect specific alterations in native endothelial cell transcripts, we made use of endothelial cell-specific RiboTag mice (Cdh5-CreERT2 mice crossed with Rpl22^HA/HA^ mice).^31,32^ EC-RiboTag mice were made hypercholesterolemic by the recombinant adeno-associated virus (AAV)-mediated hepatic overexpression of proprotein convertase subtilisin/kexin type 9 combined with a high fat diet. Thereafter, partial left carotid artery ligation was performed to induce disturbed flow and endothelial dysfunction.^50^ RiboTag-sequencing of carotid arteries 7 days after ligation (i.e. during the peak of the inflammatory response) revealed a significant upregulation of type I IFN-induced genes in the endothelium of the ligated left carotid artery (**Fig. 5A**). As our studies focused on eNOS, we next determined whether inhibiting endothelial dysfunction by preventing the inactivation of eNOS by oxidative stress could prevent the activation of type I IFN signaling. This was achieved by crossing mice expressing the redox stress-insensitive eNOS Tyr656Phe (eNOS^YF^) mutant,^51,52^ onto the apolipoprotein E-deficient (ApoE^-/-^) background. When ApoE^-/-^ mice were fed a high fat diet and subjected to partial carotid ligation, dsRNA and OAS3 accumulated in the endothelium of the ligated left carotid (**Fig. 5B**). These effects were completely abrogated in ligated left carotid arteries from ApoE-eNOS^YF^ mice, which are characterized by preserved endothelial function.^30^ Importantly, both dsRNA and the expression of OAS3 were detected in the endothelium of human carotid atherosclerotic plaques (**Fig. 5C**), indicating that the activation of type I IFN signaling characterized *in vitro* also takes place in human arteries *in vivo*.

**Fig. 5.**
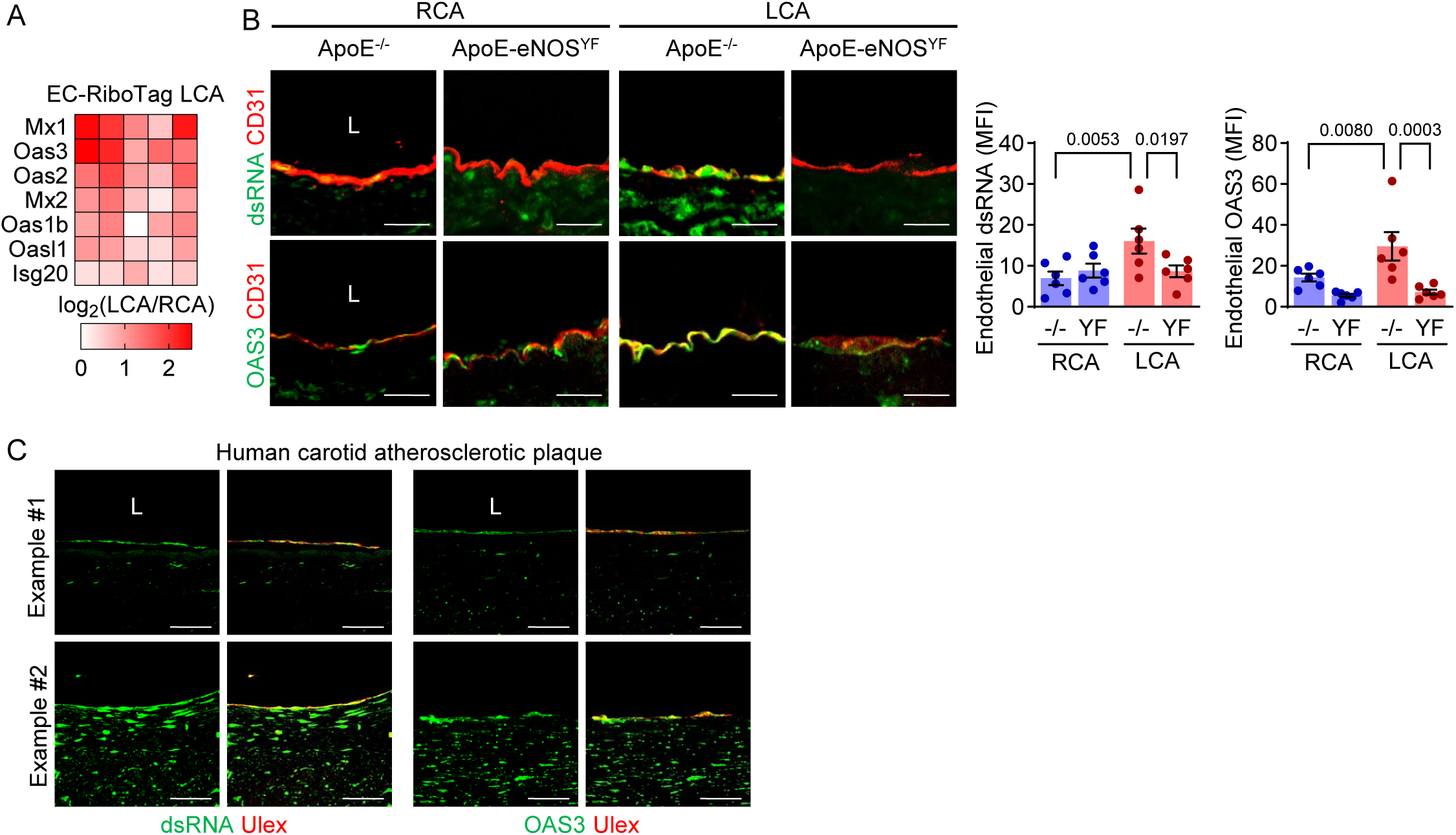
Reduced NO bioavailability induces type I IFN signaling in mice and in human. (**A**) Heatmap showing type I IFN signaling–related genes in carotid artery endothelial cells from EC-RiboTag mice following partial carotid artery ligation. LCA: left carotid artery; RCA: right carotid artery; n=5. (**B**) Confocal images showing dsRNA, OAS3 and CD31 in non-ligated right and ligated left carotid arteries from ApoE^-/-^ and ApoE-eNOS^YF^ mice 2 days after ligation; n=6 independent samples (unpaired Student’s t-test). L: lumen. Scale bar = 20 µm. (**C**) Confocal images showing dsRNA, OAS3 and Ulex (endothelial cell marker) in human carotid atherosclerotic plaques. Similar images were obtained in 11 additional patients. Scale bar = 50 µm.

## Discussion

The results of our study revealed that the eNOS-derived NO-dependent *S*-nitrosation of the A-to-I editing enzyme ADAR1 in the endothelial cell nucleus is essential to prevent the accumulation of endogenous dsRNA and the activation of a type I IFN response. We found that decreasing NO bioavailability *in vitro* by depleting eNOS or inducing endothelial cell dysfunction *in vivo*, using a combination of hypercholesterolemia and partial carotid artery ligation, triggered the accumulation of dsRNA and the induction of a series of proteins involved in IFN signaling. Importantly, the accumulation of dsRNA as well as the activation of a type I IFN response (as evidenced by OAS3 expression) could also be demonstrated in the endothelium of carotid arteries from patients with atherosclerosis.

There is a wealth of evidence demonstrating that the generation of NO by eNOS is associated with a healthy vasculature. Certainly, a genetic predisposition to enhanced NO signaling is associated with a reduced risk of developing coronary heart disease, peripheral arterial disease and stroke.^53^ Situations associated with decreased NO bioavailability, as a consequence of reduced eNOS activity or the reaction of NO with superoxide anions, on the other hand, have been linked with cardiovascular disease.^54^ Initially studies on the molecular actions of NO focused on its ability to increase cyclic GMP levels or prevent the activation of nuclear factor κB. However, it is now clear that the majority of the effects attributable to NO results from its ability to post-translationally modify proteins by *S*-nitrosation.^55,56^ The latter modification occurs preferentially where eNOS is localized,^57^ and there have been detailed analyses of *S*-nitrosated proteins in different cell compartments including mitochondria^58^ and the Golgi apparatus.^59^ The presence of eNOS in the endothelial cell nucleus has been described by several groups, as has the ability of diverse stimuli to elicit the nuclear translocation of the enzyme.^6–8^ However, little is known about the role of nuclear eNOS in regulating endothelial cell function. Given that *S*-nitrosated proteins are generally located in close vicinity to eNOS, the first step was to determine the composition of the nuclear eNOS interactome. Many of the proteins that physically associated with eNOS were involved in RNA binding, processing and splicing, which may go a long way to explaining the link between eNOS/NO signaling and alterations in endothelial gene expression. The interaction with core components of paraspeckles; small nuclear bodies involved in transcriptional regulation, was also indicative of a direct contribution of NO signaling to the regulation of gene expression.

One protein that associated with and was *S*-nitrosated by active eNOS in the endothelial cell nucleus was ADAR1. The latter protein is one of two enzymes in mammalian cells that perform A-to-I RNA editing, and is the main RNA editor in endothelial cells.^39,60^ The deamination of adenosine in dsRNA is one of the most prevalent post-transcriptional RNA modifications,^61^ and results in an I:U mismatch that is less stable than A:U base pairing and thus reduces dsRNA stability. ADAR1-mediated A-to-I editing destabilizes dsRNA structures, thereby preventing a dsRNA-triggered innate immune and inflammatory response.^60,62^ Alterations in the frequency of A-to-I editing can, therefore, result in the accumulation of dsRNA and activation of cellular responses similar to those triggered by viral infection.^63^ Indeed, in mice the endothelial cell-specific deletion or inactivation of ADAR1 leads to innate immune activation and organ dysfunction.^60^ Given that eNOS-depletion increased dsRNA levels, our next step was to determine the impact of altering NO production, and thus ADAR1 *S*-nitrosation, on A-to-I editing in endothelial cells. This identified a number of transcripts the editing of which was altered in an NO-dependent manner. The most significantly altered transcripts were EIF2AK2, VHL, SLC35F5 and the cytosolic dsRNA-binding sensor MAVS. The latter was significant inasmuch as the recognition of dsRNA by MAVS triggers the transcription of pro-inflammatory mediators as well as type I IFNs.^48,64^ In turn, type I IFNs activate a signaling cascade that increases the expression of isoforms of ADAR1 to counteract the increase in dsRNA and IFN-stimulated genes. While MAVS can be edited by the p110 and p150 isoforms of ADAR1, EIF2AK2, VHL and SLC35F5 are substrates for p150 ADAR1.^65^ Our observation that the p150 isoform was induced in eNOS-deficient cells goes a long way to explaining why the lack of NO altered rather than abolished A-to-I editing. Important for attributing a major role to the *S*-nitrosation of ADAR1 in the cellular response to eNOS depletion was the observation that both the lack of eNOS and ADAR1 induced a similar upregulation of proteins involved in type I IFN signaling. Among the type I IFN-stimulated genes robustly induced by eNOS depletion were the dsRNA-activated enzymes OAS1/2/3 and RNase L. RNase L promotes the rapid and widespread turnover of cellular RNAs, with the exception of mRNAs involved in the IFN response that are resistant to its actions.^66^ Like a typical anti-viral response, the activation of these proteins leads to cell cycle arrest and increased cell death.^49^ Both of these consequences were apparent in eNOS-deficient endothelial cells.

A clear link between endothelial dysfunction and type I IFN signaling has been established only in the context of systemic lupus erythematosus, as type I IFNs were shown to have a negative impact on eNOS expression and NO production.^67,68^ In murine models of atherosclerosis, the application of type I IFNs has been reported to promote plaque destabilization and monocyte recruitment,^69,70^ while the knockdown of the type I IFN receptor decreases atherosclerosis severity.^71^ However, these studies viewed endothelial dysfunction as a consequence of an increase in type I IFNs, rather than its cause. Our findings clearly indicate that a decreased bioavailability of nuclear eNOS-derived NO (eNOS depletion *in vitro*, endothelial dysfunction *in vivo*) leads to a marked increase in type I IFNs and their related pathways. Linking *in vitro* studies back to the *in vivo* situation is not always easy, and although it was possible to show increased dsRNA and OAS3 levels in arteries from subjects with atherosclerosis, there was no possibility to link this to NO. The situation is different in mice as OAS2 and MX2 (both type I IFN-regulated genes) were among the genes differentially regulated in ‘loss-of-function’ eNOS S1176A mice versus “gain-of-function” S1176D mice fed a high fat diet.^72^ To investigate this more closely, we made use of a previously generated eNOS mutant in which a crucial tyrosine residue (Tyr656) in the FMN binding domain that can be phosphorylated under conditions associated with oxidative stress to inactivate the enzyme was replaced with phenylalanine.^51,52^ Mice expressing this eNOS mutant (ApoE-eNOS^YF^ mice) are resistant to the development of endothelial dysfunction following hypercholesterolemia and partial carotid artery ligation.^30^ Fitting with this, we observed that the induction of atherogenesis in ApoE^-/-^ mice resulted in an increase in dsRNA and OAS3 that was more pronounced in the ligated left versus the non-ligated right carotid artery, responses that were greatly attenuated in ApoE-eNOS^YF^ mice.

Taken together, we have uncovered a novel mechanism underlying the eNOS-mediated maintenance of endothelial cell homeostasis. By regulating the *S*-nitrosation and function of ADAR1, eNOS-derived NO prevents the accumulation of dsRNA and the induction of an endothelium-driven innate immune response. In light of the link between endothelial dysfunction and the generation of type I IFNs, anti-type I IFN therapy, which showed promising results in the treatment of the cardiovascular complication associated with systemic lupus erythematosus, may be explored as novel therapeutic option in cardiovascular disease.

## Acknowledgments

X.Z. performed experiments, analyzed and interpreted data, and drafted the manuscript.

C.K. performed all bioinformatic analyses.

S.G. performed all NGS experiments and analyses.

I.W. performed the mass spectrometry analysis of the nuclear eNOS interactome.

B.G. performed the immunofluorescence staining of human carotid atherosclerotic plaques.

D.B. performed the experiments and analysis of multiplexed enhanced protein dynamic mass spectrometry.

F.D. contributed to immunofluorescence staining of mouse aorta.

N.S. and L.M. provided samples of patients with carotid atherosclerotic plaques.

C.M. contributed to the experiments and analysis of multiplexed enhanced protein dynamic mass spectrometry.

S.O. provided the EC-RiboTag mice and contributed to the interpretation of the data.

I.F. contributed major conceptual insight, interpreted the data, and wrote the manuscript.

M.S. conceived the study, designed and performed experiments, analyzed and interpreted the data and wrote the manuscript.

All the authors are indebted to Isabel Winter, Mechtild Piepenbrock-Gyamfi and Sandrine Ngaha for expert technical assistance.

Fig. 3A was created using Figdraw.

## References

1. Konst RE, Guzik TJ, Kaski J-C, Maas AHEM, Elias-Smale SE. The pathogenic role of coronary microvascular dysfunction in the setting of other cardiac or systemic conditions. Cardiovasc Res. 2020;116:817–828. doi: 10.1093/cvr/cvaa009

2. Peng HB, Libby P, Liao JK. Induction and stabilization of I kappa B alpha by nitric oxide mediates inhibition of NF-kappa B. J Biol Chem. 1995;270:14214–14219. doi: 10.1074/jbc.270.23.14214

3. Vasudevan D, Hickok JR, Bovee RC, Pham V, Mantell LL, Bahroos N, Kanabar P, Cao X-J, Maienschein-Cline M, Garcia BA, Thomas DD. Nitric oxide regulates gene expression in cancers by controlling histone posttranslational modifications. Cancer Res. 2015;75:5299–5308. doi: 10.1158/0008-5472.CAN-15-1582

4. Sessa WC, García-Cardeña G, Liu J, Keh A, Pollock JS, Bradley J, Thiru S, Braverman IM, Desai KM. The Golgi association of endothelial nitric oxide synthase is necessary for the efficient synthesis of nitric oxide. J Biol Chem. 1995;270:17641–17644. doi: 10.1074/jbc.270.30.17641

5. Shaul PW, Smart EJ, Robinson LJ, German Z, Yuhanna IS, Ying Y, Anderson RG, Michel T. Acylation targets endothelial nitric-oxide synthase to plasmalemmal caveolae. J Biol Chem. 1996;271:6518–6522. doi: 10.1074/jbc.271.11.6518

6. Feng Y, Venema VJ, Venema RC, Tsai N, Caldwell RB. VEGF induces nuclear translocation of Flk-1/KDR, endothelial nitric oxide synthase, and caveolin-1 in vascular endothelial cells. Biochem Biophys Res Commun. 1999;256:192–197. doi: 10.1006/bbrc.1998.9790

7. Nanni S, Benvenuti V, Grasselli A, Priolo C, Aiello A, Mattiussi S, Colussi C, Lirangi V, Illi B, D’Eletto M, Cianciulli AM, Gallucci M, Carli P de, Sentinelli S, Mottolese M, Carlini P, Strigari L, Finn S, Mueller E, Arcangeli G, Gaetano C, Capogrossi MC, Donnorso RP, Bacchetti S, Sacchi A, Pontecorvi A, Loda M, Farsetti A. Endothelial NOS, estrogen receptor beta, and HIFs cooperate in the activation of a prognostic transcriptional pattern in aggressive human prostate cancer. J Clin Invest. 2009;119:1093–1108. doi: 10.1172/JCI35079

8. Gobeil F, Zhu T, Brault S, Geha A, Vazquez-Tello A, Fortier A, Barbaz D, Checchin D, Hou X, Nader M, Bkaily G, Gratton J-P, Heveker N, Ribeiro-da-Silva A, Peri K, Bard H, Chorvatova A, D’Orléans-Juste P, Goetzl EJ, Chemtob S. Nitric oxide signaling via nuclearized endothelial nitric-oxide synthase modulates expression of the immediate early genes iNOS and mPGES-1. J Biol Chem. 2006;281:16058–16067. doi: 10.1074/jbc.M602219200

9. Grasselli A, Nanni S, Colussi C, Aiello A, Benvenuti V, Ragone G, Moretti F, Sacchi A, Bacchetti S, Gaetano C, Capogrossi MC, Pontecorvi A, Farsetti A. Estrogen receptor-alpha and endothelial nitric oxide synthase nuclear complex regulates transcription of human telomerase. Circ Res. 2008;103:34–42. doi: 10.1161/CIRCRESAHA.107.169037

10. Busse R, Lamontagne D. Endothelium-derived bradykinin is responsible for the increase in calcium produced by angiotensin-converting enzyme inhibitors in human endothelial cells. Naunyn Schmiedebergs Arch Pharmacol. 1991;344:126–129. doi: 10.1007/BF00167392

11. World Medical Association Declaration of Helsinki: ethical principles for medical research involving human subjects. JAMA. 2013;310:2191–2194. doi: 10.1001/jama.2013.281053

12. Fleming I, Fisslthaler B, Dixit M, Busse R. Role of PECAM-1 in the shear-stress-induced activation of Akt and the endothelial nitric oxide synthase (eNOS) in endothelial cells. J Cell Sci. 2005;118:4103–4111. doi: 10.1242/jcs.02541

13. Qian J, Zhang Q, Church JE, Stepp DW, Rudic RD, Fulton DJR. Role of local production of endothelium-derived nitric oxide on cGMP signaling and S-nitrosylation. Am J Physiol Heart Circ Physiol. 2010;298:H112–8. doi: 10.1152/ajpheart.00614.2009

14. Siragusa M, Fröhlich F, Park EJ, Schleicher M, Walther TC, Sessa WC. Stromal cell-derived factor 2 is critical for Hsp90-dependent eNOS activation. Sci Signal. 2015;8:ra81. doi: 10.1126/scisignal.aaa2819

15. Forrester MT, Foster MW, Benhar M, Stamler JS. Detection of protein S-nitrosylation with the biotin switch technique. Free Radic Biol Med. 2008;46:119–126. doi: 10.1016/j.freeradbiomed.2008.09.034

16. Bolger AM, Lohse M, Usadel B. Trimmomatic: a flexible trimmer for Illumina sequence data. Bioinformatics. 2014;30:2114–2120. doi: 10.1093/bioinformatics/btu170

17. Dobin A, Davis CA, Schlesinger F, Drenkow J, Zaleski C, Jha S, Batut P, Chaisson M, Gingeras TR. STAR: ultrafast universal RNA-seq aligner. Bioinformatics. 2013;29:15–21. doi: 10.1093/bioinformatics/bts635

18. Liao Y, Smyth GK, Shi W. featureCounts: an efficient general purpose program for assigning sequence reads to genomic features. Bioinformatics. 2014;30:923–930. doi: 10.1093/bioinformatics/btt656

19. Love MI, Huber W, Anders S. Moderated estimation of fold change and dispersion for RNA-seq data with DESeq2. Genome Biol. 2014;15:550. doi: 10.1186/s13059-014-0550-8

20. Bu D, Luo H, Huo P, Wang Z, Zhang S, He Z, Wu Y, Zhao L, Liu J, Guo J, Fang S, Cao W, Yi L, Zhao Y, Kong L. KOBAS-i: intelligent prioritization and exploratory visualization of biological functions for gene enrichment analysis. Nucleic Acids Res. 2021;49:W317–W325. doi: 10.1093/nar/gkab447

21. Piechotta M, Wyler E, Ohler U, Landthaler M, Dieterich C. JACUSA: site-specific identification of RNA editing events from replicate sequencing data. BMC Bioinformatics. 2017;18:7. doi: 10.1186/s12859-016-1432-8

22. Klann K, Tascher G, Münch C. Functional translatome proteomics reveal converging and dose-dependent regulation by mTORC1 and eIF2α. Mol Cell. 2020;77:913–925.e4. doi: 10.1016/j.molcel.2019.11.010

23. Schäfer JA, Bozkurt S, Michaelis JB, Klann K, Münch C. Global mitochondrial protein import proteomics reveal distinct regulation by translation and translocation machinery. Mol Cell. 2022;82:435–446.e7. doi: 10.1016/j.molcel.2021.11.004

24. McAlister GC, Nusinow DP, Jedrychowski MP, Wühr M, Huttlin EL, Erickson BK, Rad R, Haas W, Gygi SP. MultiNotch MS3 enables accurate, sensitive, and multiplexed detection of differential expression across cancer cell line proteomes. Anal Chem. 2014;86:7150–7158. doi: 10.1021/ac502040v

25. Klann K, Münch C. PBLMM: Peptide-based linear mixed models for differential expression analysis of shotgun proteomics data. J Cell Biochem. 2022;123:691–696. doi: 10.1002/jcb.30225

26. Szklarczyk D, Kirsch R, Koutrouli M, Nastou K, Mehryary F, Hachilif R, Gable AL, Fang T, Doncheva NT, Pyysalo S, Bork P, Jensen LJ, Mering C von. The STRING database in 2023: protein-protein association networks and functional enrichment analyses for any sequenced genome of interest. Nucleic Acids Res. 2023;51:D638–D646. doi: 10.1093/nar/gkac1000

27. Bjørklund MM, Hollensen AK, Hagensen MK, Dagnaes-Hansen F, Christoffersen C, Mikkelsen JG, Bentzon JF. Induction of atherosclerosis in mice and hamsters without germline genetic engineering. Circ Res. 2014;114:1684–1689. doi: 10.1161/CIRCRESAHA.114.302937

28. Sonntag F, Köther K, Schmidt K, Weghofer M, Raupp C, Nieto K, Kuck A, Gerlach B, Böttcher B, Müller OJ, Lux K, Hörer M, Kleinschmidt JA. The assembly-activating protein promotes capsid assembly of different adeno-associated virus serotypes. J Virol. 2011;85:12686–12697. doi: 10.1128/JVI.05359-11

29. Jungmann A, Leuchs B, Rommelaere J, Katus HA, Müller OJ. Protocol for efficient generation and characterization of adeno-associated viral vectors. Hum Gene Ther Methods. 2017;28:235–246. doi: 10.1089/hgtb.2017.192

30. Siragusa M, Thöle J, Bibli S-I, Luck B, Loot AE, Silva K de, Wittig I, Heidler J, Stingl H, Randriamboavonjy V, Kohlstedt K, Brüne B, Weigert A, Fisslthaler B, Fleming I. Nitric oxide maintains endothelial redox homeostasis through PKM2 inhibition. EMBO J. 2019;38:e100938. doi: 10.15252/embj.2018100938

31. Pitulescu ME, Schmidt I, Benedito R, Adams RH. Inducible gene targeting in the neonatal vasculature and analysis of retinal angiogenesis in mice. Nat Protoc. 2010;5:1518–1534. doi: 10.1038/nprot.2010.113

32. Sanz E, Yang L, Su T, Morris DR, McKnight GS, Amieux PS. Cell-type-specific isolation of ribosome-associated mRNA from complex tissues. Proc Natl Acad Sci USA. 2009;106:13939–13944. doi: 10.1073/pnas.0907143106

33. Kumar S, Kang D-W, Rezvan A, Jo H. Accelerated atherosclerosis development in C57Bl6 mice by overexpressing AAV-mediated PCSK9 and partial carotid ligation. Lab Invest. 2017;97:935–945. doi: 10.1038/labinvest.2017.47

34. Perez-Riverol Y, Csordas A, Bai J, Bernal-Llinares M, Hewapathirana S, Kundu DJ, Inuganti A, Griss J, Mayer G, Eisenacher M, Pérez E, Uszkoreit J, Pfeuffer J, Sachsenberg T, Yilmaz S, Tiwary S, Cox J, Audain E, Walzer M, Jarnuczak AF, Ternent T, Brazma A, Vizcaíno JA. The PRIDE database and related tools and resources in 2019: improving support for quantification data. Nucleic Acids Res. 2019;47:D442–D450. doi: 10.1093/nar/gky1106

35. Mnatsakanyan R, Markoutsa S, Walbrunn K, Roos A, Verhelst SHL, Zahedi RP. Proteome-wide detection of S-nitrosylation targets and motifs using bioorthogonal cleavable-linker-based enrichment and switch technique. Nat Commun. 2019;10:2195. doi: 10.1038/s41467-019-10182-4

36. Ben-Lulu S, Ziv T, Admon A, Weisman-Shomer P, Benhar M. A substrate trapping approach identifies proteins regulated by reversible S-nitrosylation. Mol Cell Proteomics. 2014;13:2573–2583. doi: 10.1074/mcp.M114.038166

37. Wu C, Jain MR, Li Q, Oka S-I, Li W, Kong A-NT, Nagarajan N, Sadoshima J, Simmons WJ, Li H. Identification of novel nuclear targets of human thioredoxin 1. Mol Cell Proteomics. 2014;13:3507–3518. doi: 10.1074/mcp.M114.040931

38. Chung HS, Murray CI, Venkatraman V, Crowgey EL, Rainer PP, Cole RN, Bomgarden RD, Rogers JC, Balkan W, Hare JM, Kass DA, van Eyk JE. Dual labeling biotin switch assay to reduce bias derived from different cysteine subpopulations: a method to maximize S-nitrosylation detection. Circ Res. 2015;117:846–857. doi: 10.1161/CIRCRESAHA.115.307336

39. Stellos K, Gatsiou A, Stamatelopoulos K, Perisic Matic L, John D, Lunella FF, Jaé N, Rossbach O, Amrhein C, Sigala F, Boon RA, Fürtig B, Manavski Y, You X, Uchida S, Keller T, Boeckel J-N, Franco-Cereceda A, Maegdefessel L, Chen W, Schwalbe H, Bindereif A, Eriksson P, Hedin U, Zeiher AM, Dimmeler S. Adenosine-to-inosine RNA editing controls cathepsin S expression in atherosclerosis by enabling HuR-mediated post-transcriptional regulation. Nat Med. 2016;22:1140–1150. doi: 10.1038/nm.4172

40. Brümmer A, Yang Y, Chan TW, Xiao X. Structure-mediated modulation of mRNA abundance by A-to-I editing. Nat Commun. 2017;8:1255. doi: 10.1038/s41467-017-01459-7

41. Lamers MM, van den Hoogen BG, Haagmans BL. ADAR1: “editor-in-chief” of cytoplasmic innate immunity. Front Immunol. 2019;10:1763. doi: 10.3389/fimmu.2019.01763

42. Bavelloni A, Focaccia E, Piazzi M, Raffini M, Cesarini V, Tomaselli S, Orsini A, Ratti S, Faenza I, Cocco L, Gallo A, Blalock WL. AKT-dependent phosphorylation of the adenosine deaminases ADAR-1 and -2 inhibits deaminase activity. FASEB J. 2019;33:9044–9061. doi: 10.1096/fj.201800490RR

43. Desterro JMP, Keegan LP, Jaffray E, Hay RT, O’Connell MA, Carmo-Fonseca M. SUMO-1 modification alters ADAR1 editing activity. Mol Biol Cell. 2005;16:5115–5126. doi: 10.1091/mbc.e05-06-0536

44. Sun Q, Sun L, Liu H-H, Chen X, Seth RB, Forman J, Chen ZJ. The specific and essential role of MAVS in antiviral innate immune responses. Immunity. 2006;24:633– 642. doi: 10.1016/j.immuni.2006.04.004

45. Hou F, Sun L, Zheng H, Skaug B, Jiang Q-X, Chen ZJ. MAVS forms functional prion-like aggregates to activate and propagate antiviral innate immune response. Cell. 2011;146:448–461. doi: 10.1016/j.cell.2011.06.041

46. Jain M, Jantsch MF, Licht K. The editor’s I on disease development. Trends Genet. 2019;35:903–913. doi: 10.1016/j.tig.2019.09.004

47. Uggenti C, Lepelley A, Crow YJ. Self-awareness: nucleic acid-driven inflammation and the type I interferonopathies. Annu Rev Immunol. 2019;37:247–267. doi: 10.1146/annurev-immunol-042718-041257

48. Chen YG, Hur S. Cellular origins of dsRNA, their recognition and consequences. Nat Rev Mol Cell Biol. 2022;23:286–301. doi: 10.1038/s41580-021-00430-1

49. Li Y, Banerjee S, Goldstein SA, Dong B, Gaughan C, Rath S, Donovan J, Korennykh A, Silverman RH, Weiss SR. Ribonuclease L mediates the cell-lethal phenotype of double-stranded RNA editing enzyme ADAR1 deficiency in a human cell line. Elife. 2017;6. doi: 10.7554/eLife.25687

50. Nam D, Ni C-W, Rezvan A, Suo J, Budzyn K, Llanos A, Harrison D, Giddens D, Jo H. Partial carotid ligation is a model of acutely induced disturbed flow, leading to rapid endothelial dysfunction and atherosclerosis. Am J Physiol Heart Circ Physiol. 2009;297:H1535–43. doi: 10.1152/ajpheart.00510.2009

51. Fisslthaler B, Loot AE, Mohamed A, Busse R, Fleming I. Inhibition of endothelial nitric oxide synthase activity by proline-rich tyrosine kinase 2 in response to fluid shear stress and insulin. Circ Res. 2008;102:1520–1528. doi: 10.1161/CIRCRESAHA.108.172072

52. Loot AE, Schreiber JG, Fisslthaler B, Fleming I. Angiotensin II impairs endothelial function via tyrosine phosphorylation of the endothelial nitric oxide synthase. J Exp Med. 2009;206:2889–2896. doi: 10.1084/jem.20090449

53. Emdin CA, Khera AV, Klarin D, Natarajan P, Zekavat SM, Nomura A, Haas M, Aragam K, Ardissino D, Wilson JG, Schunkert H, McPherson R, Watkins H, Elosua R, Bown MJ, Samani NJ, Baber U, Erdmann J, Gormley P, Palotie A, Stitziel NO, Gupta N, Danesh J, Saleheen D, Gabriel S, Kathiresan S. Phenotypic consequences of a genetic predisposition to enhanced nitric oxide signaling. Circulation. 2018;137:222–232. doi: 10.1161/CIRCULATIONAHA.117.028021

54. Förstermann U, Xia N, Li H. Roles of vascular oxidative stress and nitric oxide in the pathogenesis of atherosclerosis. Circ Res. 2017;120:713–735. doi: 10.1161/CIRCRESAHA.116.309326

55. Haldar SM, Stamler JS. S-nitrosylation: integrator of cardiovascular performance and oxygen delivery. J Clin Invest. 2013;123:101–110. doi: 10.1172/JCI62854

56. Greco TM, Hodara R, Parastatidis I, Heijnen HFG, Dennehy MK, Liebler DC, Ischiropoulos H. Identification of S-nitrosylation motifs by site-specific mapping of the S-nitrosocysteine proteome in human vascular smooth muscle cells. Proc Natl Acad Sci U S A. 2006;103:7420–7425. doi: 10.1073/pnas.0600729103

57. Iwakiri Y, Satoh A, Chatterjee S, Toomre DK, Chalouni CM, Fulton D, Groszmann RJ, Shah VH, Sessa WC. Nitric oxide synthase generates nitric oxide locally to regulate compartmentalized protein S-nitrosylation and protein trafficking. Proc Natl Acad Sci U S A. 2006;103:19777–19782. doi: 10.1073/pnas.0605907103

58. Boulghobra D, Dubois M, Alpha-Bazin B, Coste F, Olmos M, Gayrard S, Bornard I, Meyer G, Gaillard J-C, Armengaud J, Reboul C. Increased protein S-nitrosylation in mitochondria: a key mechanism of exercise-induced cardioprotection. Basic Res Cardiol. 2021;116:66. doi: 10.1007/s00395-021-00906-3

59. Sangwung P, Greco TM, Wang Y, Ischiropoulos H, Sessa WC, Iwakiri Y. Proteomic identification of S-nitrosylated Golgi proteins: new insights into endothelial cell regulation by eNOS-derived NO. PLoS One. 2012;7:e31564. doi: 10.1371/journal.pone.0031564

60. Guo X, Liu S, Yan R, Nguyen V, Zenati M, Billiar TR, Wang Q. ADAR1 RNA editing regulates endothelial cell functions via the MDA-5 RNA sensing signaling pathway. Life Sci Alliance. 2021;5. doi: 10.26508/lsa.202101191

61. Walkley CR, Li JB. Rewriting the transcriptome: adenosine-to-inosine RNA editing by ADARs. Genome Biol. 2017;18:205. doi: 10.1186/s13059-017-1347-3

62. Garcia-Gonzalez C, Dieterich C, Maroli G, Wiesnet M, Wietelmann A, Li X, Yuan X, Graumann J, Stellos K, Kubin T, Schneider A, Braun T. ADAR1 prevents autoinflammatory processes in the heart mediated by IRF7. Circ Res. 2022;131:580–597. doi: 10.1161/CIRCRESAHA.122.320839

63. Rivera M, Zhang H, Pham J, Isquith J, Zhou QJ, Balaian L, Sasik R, Enlund S, Mark A, Ma W, Holm F, Fisch KM, Kuo DJ, Jamieson C, Jiang Q. Malignant A-to-I RNA editing by ADAR1 drives T cell acute lymphoblastic leukemia relapse via attenuating dsRNA sensing. Cell Rep. 2024;43:113704. doi: 10.1016/j.celrep.2024.113704

64. Dias Junior AG, Sampaio NG, Rehwinkel J. A balancing act: MDA5 in antiviral immunity and autoinflammation. Trends Microbiol. 2018;27:75–85. doi: 10.1016/j.tim.2018.08.007

65. Sun T, Yu Y, Wu X, Acevedo A, Luo J-D, Wang J, Schneider WM, Hurwitz B, Rosenberg BR, Chung H, Rice CM. Decoupling expression and editing preferences of ADAR1 p150 and p110 isoforms. Proc Natl Acad Sci U S A. 2021;118. doi: 10.1073/pnas.2021757118

66. Burke JM, Moon SL, Matheny T, Parker R. RNase L reprograms translation by widespread mRNA turnover escaped by antiviral mRNAs. Mol Cell. 2019;75:1203–1217.e5. doi: 10.1016/j.molcel.2019.07.029

67. Buie JJ, Renaud LL, Muise-Helmericks R, Oates JC. IFN-α negatively regulates the expression of endothelial nitric oxide synthase and nitric oxide production: implications for systemic Lupus Erythematosus. J Immunol. 2017;199:1979–1988. doi: 10.4049/jimmunol.1600108

68. Jia H, Thelwell C, Dilger P, Bird C, Daniels S, Wadhwa M. Endothelial cell functions impaired by interferon in vitro: Insights into the molecular mechanism of thrombotic microangiopathy associated with interferon therapy. Thromb Res. 2018;163:105–116. doi: 10.1016/j.thromres.2018.01.039

69. Goossens P, Gijbels MJJ, Zernecke A, Eijgelaar W, Vergouwe MN, van der Made I, Vanderlocht J, Beckers L, Buurman WA, Daemen MJAP, Kalinke U, Weber C, Lutgens E, Winther MPJ de. Myeloid type I interferon signaling promotes atherosclerosis by stimulating macrophage recruitment to lesions. Cell Metab. 2010;12:142–153. doi: 10.1016/j.cmet.2010.06.008

70. Venkatesh D, Ernandez T, Rosetti F, Batal I, Cullere X, Luscinskas FW, Zhang Y, Stavrakis G, García-Cardeña G, Horwitz BH, Mayadas TN. Endothelial TNF receptor 2 induces IRF1 transcription factor-dependent interferon-β autocrine signaling to promote monocyte recruitment. Immunity. 2013;38:1025–1037. doi: 10.1016/j.immuni.2013.01.012

71. Thacker SG, Zhao W, Smith CK, Luo W, Wang H, Vivekanandan-Giri A, Rabquer BJ, Koch AE, Pennathur S, Davidson A, Eitzman DT, Kaplan MJ. Type I interferons modulate vascular function, repair, thrombosis, and plaque progression in murine models of lupus and atherosclerosis. Arthritis Rheum. 2012;64:2975–2985. doi: 10.1002/art.34504

72. Nguyen TD, Rahman N-T, Sessa WC, Lee MY. Endothelial nitric oxide synthase (eNOS) S1176 phosphorylation status governs atherosclerotic lesion formation. Front Cardiovasc Med. 2023;10:1279868. doi: 10.3389/fcvm.2023.1279868

